# Extracellular matrix remodeling through endocytosis and resurfacing of Tenascin-R

**DOI:** 10.1101/2020.11.11.377515

**Authors:** Tal M. Dankovich, Rahul Kaushik, Gabriel Cassinelli Petersen, Philipp Emanuel Giro, Hannah Abdul Hadi, Guobin Bao, Sabine Beuermann, Benjamin H. Cooper, Alexander Dityatev, Silvio O. Rizzoli

## Abstract

The brain extracellular matrix (ECM) assembles around neurons and synapses, and is thought to change only rarely, through proteolysis and renewed protein synthesis. We report here an alternative ECM remodeling mechanism, based on the recycling of ECM molecules. We found that a key ECM protein, Tenascin-R, is frequently endocytosed, and later resurfaces, preferentially near synapses. The TNR molecules complete this cycle within ∼3 days, in an activity-dependent fashion.

## Main text

The extracellular matrix (ECM) molecules of the brain form lattices that enwrap neurons and fill perisynaptic spaces ^1^. These structures appear to be particularly durable, owing to the exceptional longevity of the ECM molecules ^2,3^, and they are deemed to stabilize neurons and synapses ^4,5^. Nevertheless, the ECM must remain alterable to allow for the structural changes that occur in response to activity and plasticity. Such changes have long been thought to be infrequent in the adult brain, but this notion has been recently challenged by a series of super-resolution imaging studies. Synapses were found to change shape and location continually, on a time scale of minutes, both in acute brain slices ^6^ and in the adult brain ^7,8^. These findings therefore suggest that the ECM may be remodeled relatively frequently.

This notion is difficult to accommodate with the best-known mechanisms for ECM remodeling, which involve ECM cleavage by proteolytic enzymes such as matrix metalloproteinases, followed by the secretion and incorporation of newly-synthesized ECM molecules ^9,10^. An alternative solution to ECM remodeling seems therefore necessary, through mechanisms that reuse existing molecules, rather than relying on *de novo* synthesis. To search for such a mechanism, we focused on Tenascin-R (TNR), a matrix glycoprotein that is predominantly expressed in the central nervous system ^11^. TNR is highly enriched in perineuronal nets (PNNs), a condensed ECM lattice surrounding a subset of inhibitory interneurons, and is essential for PNN formation. TNR is also expressed in the more diffuse perisynaptic ECM associated with both inhibitory and excitatory synapses on a broad range of neuronal cell types ^12^ (see Extended Data Fig. 1a-b).

To test whether ECM molecules can indeed be reused, we first used a classical biotinylation-based assay that has been instrumental in determining the reuse (recycling) of neuronal surface proteins, and especially of neurotransmitter receptors ^13^. We employed the same system used previously for determining the recycling of neurotransmitter receptors, rat cultured hippocampal neurons. We treated the neurons with a cleaveable, membrane-impermeable biotin derivative, to tag all proteins at the cell surface. We allowed the neurons to internalize molecules for 6 hours, and we then stripped biotin from the cell surface, thus leaving this label only on the endocytosed proteins (Fig. 1a). To measure the potential recycling of the internalized molecules, we incubated the neurons a further 18 hours, to allow for protein resurfacing, and then performed a second round of biotin stripping (Fig. 1a). We collected the biotin-tagged proteins on streptavidin beads, and we then tested the recycling of TNR and other proteins by immunolabeling the beads and imaging them in confocal microscopy (Fig. 1b). This is a particularly sensitive technique, enabling the detection of proteins in minute sample volumes ^14^. We found that TNR is indeed endocytosed during the initial 6 hours of incubation, and that a significant proportion of the molecules resurfaces during the subsequent 18 hours. This behavior was similar to that of a positive control, the synaptic vesicle protein synaptotagmin 1, Syt1, which participates in the well-known recycling of synaptic vesicles (Fig. 1c). The loss of biotinylated TNR cannot be ascribed to its degradation, since this molecule is extremely stable (Extended Data Fig. 1c-d), and since we blocked lysosomal degradation during these experiments.

**Fig. 1:**
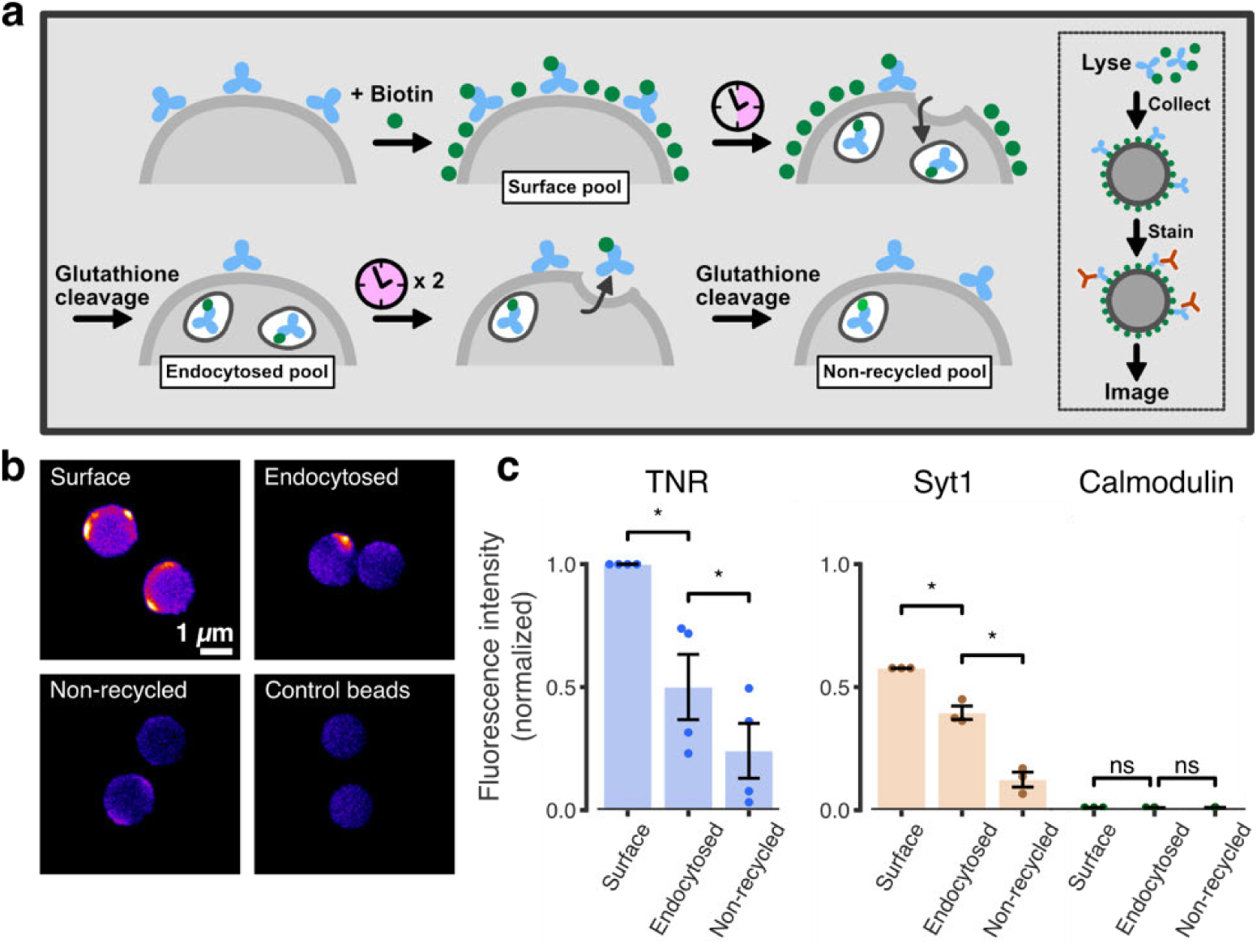
TNR molecules endocytose and subsequently resurface in neurons. **a**, We assessed the recycling of TNR molecules biochemically using a cell-surface biotinylation assay. We incubated live neurons with the reversible membrane-impermeable biotin derivative sulfo-NHS-S-S-biotin, to label all cell-surface proteins. We then incubated the neurons at 37°C for 6 hours, to allow for internalization, and the remaining cell-surface proteins were stripped of their biotin labels by incubation with glutathione. The internalized proteins, which are protected from the glutathione cleavage, constitute the ‘endocytosed pool’. The neurons were incubated at 37°C a further 18 hours, to allow for recycling back to the cell surface, and then exposed to a second round of glutathione cleavage. The internalized proteins following the second round of stripping constitute the ‘non-recycled pool’. Neuronal lysates from these different experiments, containing biotin-labeled proteins representing the surface, endocytosed or non-recycled molecules, were then collected on streptavidin-coupled beads, and were immunostained with TNR antibodies, together with fluorescent-labeled anti-mouse secondary nanobodies. The stained beads were then immobilized on glass slides and imaged with confocal microscopy, to quantify the TNR amounts. **b**, Example images of beads collecting the surface, endocytosed and non-recycled TNR pools (top-left, top-right and bottom-left panels respectively). Bottom-right panel: control beads, incubated with fluorescent-labeled secondary nanobodies, in the absence of TNR antibodies. Scale bar = 1 µm. **c**, The graphs show the fluorescence intensity of the beads, normalized to the mean of the ‘surface’ condition in the corresponding experiment. An analysis shows that a substantial fraction of surface TNR is endocytosed within 6 hours after labeling, and that many of the molecules recycle to the surface within 24 hours (n = 4 individual experiments, at least 100 beads imaged per experiment; repeated-measures one-way ANOVA: F_1.153, 3.458_ = 28.29, *p*\*\* = 0.009, followed by Holm-Sidak multiple comparisons test: *p*\* = 0.032 and *p*\* = 0.021 for the comparisons between the ‘surface’ and ‘endocytosed’, and ‘endocytosed’ and ‘non-recycled’ conditions, respectively). The assay was repeated for the synaptic vesicle protein Syt1, which is well known to undergo recycling, and therefore serves as a positive control, and the intracellular protein calmodulin, as a negative control. The plots for Syt1 and calmodulin were scaled by the ratio between the mean surface intensity for these proteins and that of TNR. As expected, a recycling dynamic was observed for Syt1, whereas no significant change was observed for calmodulin (n = 3 individual experiments, at least 50 beads imaged per experiment, RM one-way ANOVA: F_1.007, 2.014_ = 62.98, \**p* = 0.015; F_1, 2_ = 0.016, *p* = 0.912, followed by Holm-Sidak multiple comparisons test: *p*\* = 0.044 and *p*\* = 0.044; *p* = 0.933 and *p* = 0.993 for Syt1 and calmodulin, respectively). All data represent the mean ± SEM, with dots indicating the individual experiments.

To verify these findings by an imaging assay, we turned to a knock-out validated TNR antibody (Extended Data Fig. 2a-b). Application of the antibody to neuronal cultures did not appear to cause any changes in their behavior, as was verified by electrophysiological measurements (Extended Data Fig. 2c-d). We applied fluorophore-conjugated TNR antibodies to the cultures, and imaged them at 37°C, for several hours. We observed the accumulation of TNR antibodies in the neuronal somas, indicative of endocytosis (Extended Data Fig. 3). However, many other cellular regions remained virtually unchanged, which suggests that not all TNR molecules are dynamic.

To focus on the dynamic TNR molecules, we relied on an assay used extensively for synaptic vesicle proteins, a ‘blocking-labeling’ assay (see ^15^ and references therein). The surface TNR epitopes were blocked using unconjugated antibodies. Fluorophore-conjugated TNR antibodies were then applied at different intervals, to label newly-emerged epitopes (Fig. 2a). This assay enabled us to detect a slow but steady appearance of such epitopes (Fig. 2b,c). A super-resolution investigation using STED microscopy revealed that these TNR molecules were enriched near synapses (Fig. 2d), where they also emerged more rapidly than for the cell as a whole (compare Fig. 2c and Fig. 2e). Importantly, their appearance was potentiated by enhancing neuronal activity using the GABA_A_ blocker bicuculline, and was inhibited by reducing neuronal activity using the glutamate receptor blockers CNQX/AP5 (Fig. 2f). Moreover, the newly-emerged epitopes were found in higher amounts at synapses with larger active vesicle pools and at synapses with larger postsynaptic spines (Extended Data Fig. 4). This suggests that the emergence of these TNR molecules on the surface correlates to synaptic weight.

**Fig. 2:**
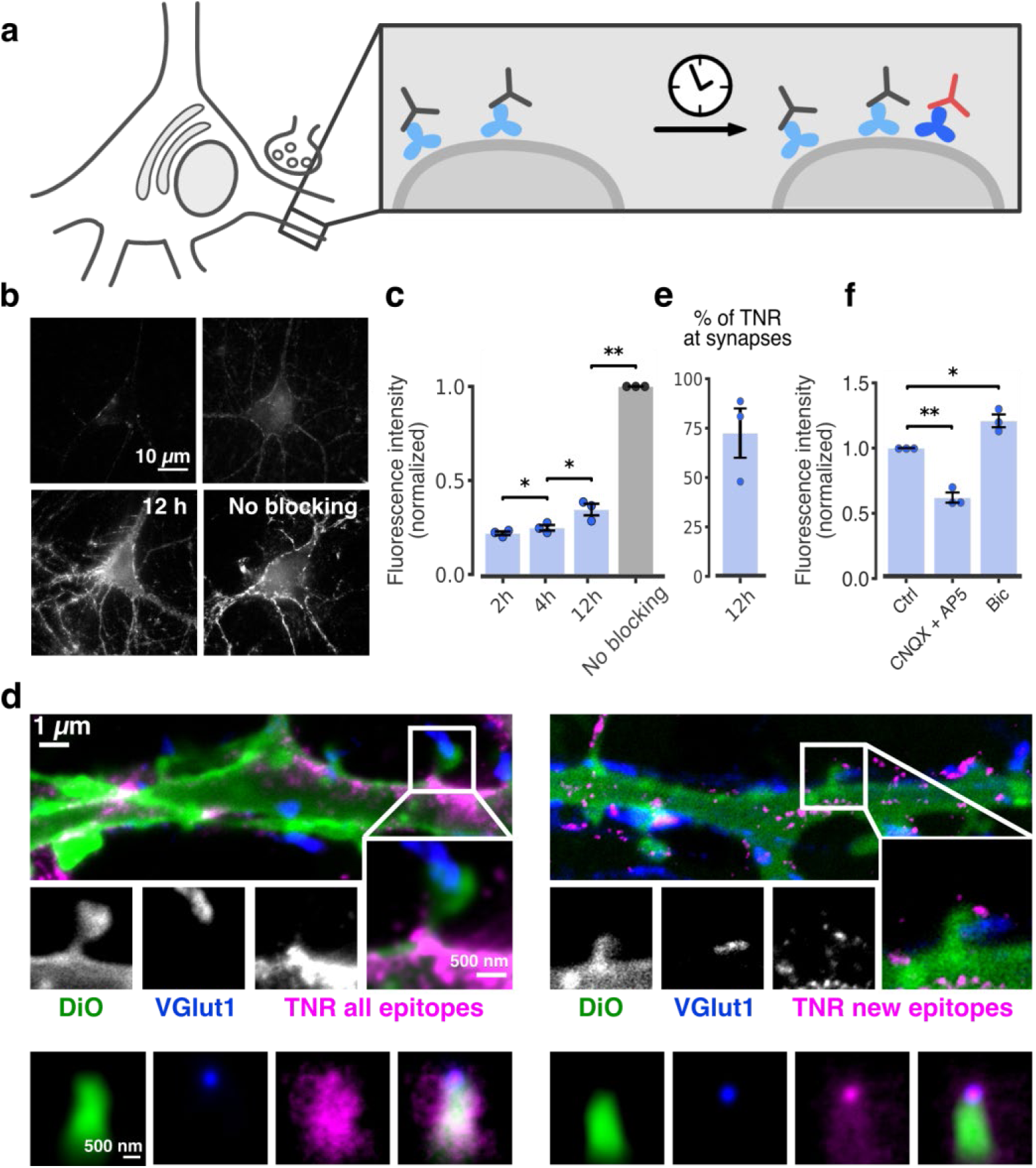
Dynamic TNR molecules emerge at synapses, in an activity-dependent fashion. **a**, To monitor dynamic TNR molecules, we first blocked the surface TNR epitopes (light blue) by incubating live neurons with non-fluorescent antibodies (gray). After a time interval (2-12 hours) we added fluorophore-conjugated TNR antibodies (red) to reveal the newly-emerged TNR epitopes (dark blue). These molecules can be then detected in imaging experiments. **b**, Newly-emerged TNR epitopes were revealed 2, 4 and 12 hours after surface epitope blocking, imaged by epifluorescence microscopy. Scale bar = 10 µm. **c**, The graph shows the fluorescence intensity normalized to that of non-blocked neurons, in which all TNR epitopes in the ECM were labeled. To avoid issues such as saturation or out-of-focus signals, the analysis concentrated on the neurites, and avoided the cell bodies. Approximately 34% of the TNR epitopes exchange (*i*.*e*. are replaced by newly-emerged TNR epitopes) within 12 hours (n = 3 independent experiments, at least 10 neurons imaged per data point; repeated measures one-way ANOVA: F_1.089, 2.179_ = 790.8, \*\*\**p <* 0.001, followed by Fisher’s LSD test: \**p =* 0.041, \**p =* 0.032 and \*\**p =* 0.002 for the comparisons between ‘2 h’ and ‘4 h’ ‘4 h’ and ‘12 h’, and ‘12 h’ and ‘no blocking’ conditions respectively). **d**, To analyze the location of the newly-emerged epitopes, we analyzed these stainings using STED microscopy. All TNR epitopes from the ECM (left panel) or only epitopes emerged within 12 hours after blocking (right panel) were revealed using Atto647N-conjugated TNR antibodies (magenta). Neuronal plasma membranes were labeled using DiO (green), and presynaptic boutons were identified by immunostaining for the synaptic vesicle marker VGlut1 (blue). We used a sparse DiO labeling, which reveals all neurites from individual neurons, but not all neurites in the cultures. The images show dendrites of DiO-labeled neurons, which form synapses with VGlut1-positive boutons from DiO-free axons. TNR and VGlut1 are imaged in 2-color STED, DiO in confocal microscopy. Scale bars: 1 µm for the initial images, 500 nm for the insets. As expected, fewer TNR epitopes are detected after blocking. A high proportion is found close to presynaptic boutons. Bottom panels: to analyze the epitope localization, we averaged several hundred synapses, from three independent experiments, by centering synapse images on the VGlut1 puncta and orienting the dendritic DiO signals vertically. The resulting distribution for all TNR epitopes covers the entire bouton-dendrite area (left panel). When the same procedure is performed for the newly-emerged TNR epitopes, a substantial enrichment in the bouton region is found (right panel). A statistical analysis comparing the enrichment in the bouton region indicated that this parameter is substantially higher for the newly-emerged TNR epitopes. This was found both when analyzing DiO-labeled dendrites and axons (n = >100 synapses analyzed for each experiment; repeated measures ANOVA on log-transformed data: F_1.977, 3.954_ = 24.13, \*\**p =* 0.006, followed by Fisher’s LSD test: \**p =* 0.024 and \**p* = 0.036 for the comparison of ‘new epitopes’ to ‘all epitopes’ at distance = 0, for dendrites and axons respectively). **e**, We measured the amount of newly-emerged TNR at synapses at 12 hours after blocking, by estimating the mean fluorescence intensity in VGlut1-positive pixels, as visualized in panel d. 75% of the epitopes at synapses exchange in 12 hours (n = 3 independent experiments, with >100 synapses analyzed for each experiment). **f**, We labeled newly-emerged TNR epitopes 12 hours after blocking, comparing control cultures with cultures in which network activity was enhanced by inhibiting GABA_A_ receptors using bicuculline (40 µM), or cultures silenced by inhibiting AMPA and NMDA receptors with CNQX (10 µM) and AP5 (50 µM). The cultures were then imaged in epifluorescence microscopy, as in panel b. The graphs show the mean fluorescence intensity normalized to the to the corresponding control (DMSO-treated) condition. A comparison of the intensities indicates that the treatments had a significant effect, with bicuculline increasing and CNQX+AP5 reducing the amount of newly-emerged TNR epitopes (n = 3 comparison experiments per treatment condition, at least 10 neurons imaged per data point; one-way ANOVA on rank-transformed data: F_2, 6_ = 42, \*\*\**p* < 0.001, followed by the Dunn multiple comparisons test: \*\**p =* 0.003 and \**p* = 0.042, for the comparison of ‘ctrl’ to ‘CNQX+AP5’ and ‘bic’, respectively). As for panel c, neurites were analyzed here, avoiding cell bodies. All data represent the mean ± SEM, with dots indicating the individual experiments.

Overall, these experiments suggest that this assay is able to detect TNR molecules appearing on the surface, from an intracellular TNR population. However, new TNR epitopes could also emerge through the un-binding of unconjugated antibodies from their epitopes, which would allow the fluorophore-conjugated ones to take their place. In control experiments, we found no evidence for such un-binding, either from the surface of fixed cells at 37°C (Extended Data Fig. 5a-b) or in live cells at 4°C (Extended Data Fig. 5c-f). A second possible source for such epitopes would be a pre-existing population of molecules that were present on the cell surface, but were previously unavailable to antibody binding, due to effects such as steric hindrance. Such molecules would be revealed by antibodies when the steric hindrance is eliminated by treatments that change the neuronal surface profoundly. To test this, we subjected the neurons to treatments that modify the membrane proteins (aldehyde treatment), or that remove glycan chains (chondroitinase ABC). We found that such treatments cause no changes in TNR epitope availability (Extended Data Fig. 5g). A third potential source of new TNR epitopes is the proteolytic cleavage of pre-existing surface TNR molecules or their binding partners, which would reduce steric hindrance and make TNR epitopes available for antibody binding. We found no evidence for this, since the TNR epitopes appeared in the same fashion after blocking the activity of matrix metalloproteinases that might cleave existing ECM (Extended Data Fig. 5h).

We therefore conclude that neurons contain a dynamic pool of TNR molecules, which appear on the surface preferentially near synapses (Fig. 2). We imaged these epitopes in living cells, and found that after surfacing they are endocytosed on a time scale of hours (Fig. 3a). To visualize the location of the internalized molecules in neurites, we allowed endocytosis to proceed for several hours, and then eliminated all surface molecules by a proteinase K treatment (Fig. 3b). We found internalized TNR to be present in both axons and dendrites (Fig. 3b). To then verify whether these molecules resurface, we designed an assay in which we tested the amounts of antibody-labeled TNR present on the surface at different time points (Fig. 3c). Immediately after antibody labeling of TNR, the antibodies are found mostly on the surface, as expected, and neurites are fully visible. A day or two later, the antibodies are no longer on the surface, as they have been endocytosed, and, since many of the organelles have already reached the cell body (Fig. 3a), neurites are virtually invisible. Remarkably, this situation changes at 3 days after labeling, and a high proportion of the antibodies are again on the surface, especially in neurites (Fig. 3c). These antibodies are later again endocytosed, and will return to the surface after another 3 days (Fig. 3c).

**Fig. 3:**
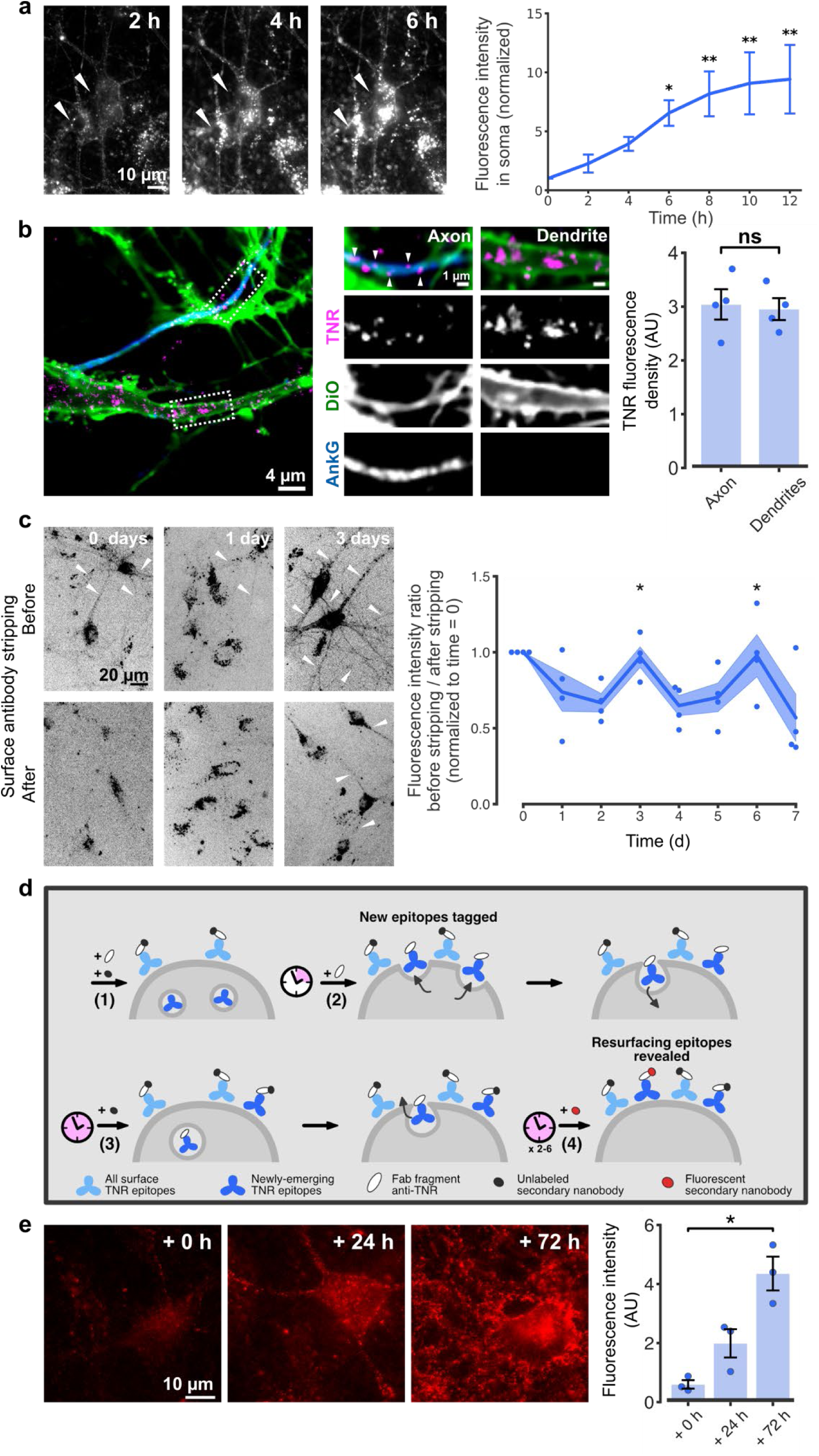
Dynamic TNR molecules are endocytosed in neurons over hours, and recycle with a periodicity of ∼3 days. **a**, The newly-emerged TNR epitopes were labeled using fluorophore-conjugated antibodies (at 4 hours after blocking), and were monitored by live epifluorescence imaging over 12 hours. A strong internalization of TNR epitopes is evident, as fluorescence accumulates in the cell somas (several indicated by arrowheads). Scale bar = 10 μm. The plot shows the mean fluorescence intensity in the cell somas, normalized to the t_0_ timepoint. A significant increase in the signal over time confirms the observation that the TNR molecules are endocytosed (n = 5 independent experiments, Friedman test: χ^2^_6_ = 25.46, \*\*\**p* < 0.001, followed by the Dunn multiple comparisons test: \**p* = 0.033, \*\**p* = 0.005, \*\**p* = 0.005, \*\**p* = 0.002 for 6, 8, 10 and 12 hours respectively). Data represent the mean ± SEM. **b**, The presence of internalized TNR molecules in cell bodies is evident in panel a, but the presence of such molecules in neurites is difficult to analyze without further experimental steps. To address this, we labeled newly-emerged TNR epitopes as in a (magenta), and allowed them to internalize for 12 hours. The surface molecules were then removed by incubation with proteinase K, and the remaining TNR signal density, which corresponds to *bona fide* internalized molecules, was measured in both dendritic and axonal processes. The neurites were identified in DiO stainings (green). An additional immunostaining for AnkyrinG (blue), which labels the axon initial segment, was used to identify the axons. Processes that did not overlap with the AnkyrinG staining, and showed evident postsynaptic spines, were classified as dendrites. Scale bar = 4 μm. The images were taken with confocal microscopy. An analysis of the average TNR intensity (equivalent to signal density) indicates that internalized TNR epitopes are equally present in dendrites and axons. As observed in the zoom panels, the larger volume of the particular dendrites implies that substantially more TNR-positive objects were seen there. However, the overall signal density (which takes the difference in volume into account) is similar for dendrites and axons. Statistical significance was evaluated using a paired *t*-test (n = 4 experiments, at least 10 neurons imaged per data point, t = 0.741, *p =* 0.513). Data represent the mean ± SEM, with dots indicating the individual experiments. **c**, To test whether the internalized TNR molecules recycle to the ECM, we labeled them as in the other panels, and we then measured the fraction present on the surface after different time intervals. To determine this, we imaged the neurons in epifluorescence before and after stripping the surface molecules using proteinase K. As for the epifluorescence images in Fig. 2, we analyzed the signals in the neurites, and avoided the cell bodies. At day 0 (immediately after labeling), the stripping procedure strongly reduced the staining, causing neurites to become virtually invisible after stripping. The effect was less pronounced at 1 day after staining. At this time point only few neurites were evident even before stripping, so this treatment did not cause a strong change. The effect became again evident at 3 days after staining. Neurites were again abundantly visible, and again most disappeared after stripping, indicating that a high proportion of TNR molecules returned to the surface at 3 days after labeling. We quantified this by reporting the fluorescence ratio between the images taken before and after stripping (normalized to the day 0 time point). The graph indicates that a similar peak of TNR return to the surface is also evident at 6 days after labeling (∼3 day periodicity). The amounts stripped at days 3 and 6 are significantly higher than at days 1 and 2, or 5 and 7 (n = 4 independent experiments). Statistical significance was evaluated using the Kruskal-Wallis test followed by Fisher’s LSD (Days 2, 3, 4: H_2_ = 8.29, \**p* = 0.016, day 3 vs. day 2: \**p* = 0.046, day 3 vs. day 4: \*\**p* = 0.005; Days 4, 5, 6: H_2_ = 6.74, \**p* = 0.036, day 6 vs. day 5: \**p* = 0.022, day 6 vs. day 7: \**p* = 0.028). Scale bar = 20 μm. Data represent the mean (lines) ± SEM (shaded regions); dots indicate the individual experiments. **d**, To better identify the actively recycling TNR molecules, we developed an assay to specifically label only those molecules that had completed a full endocytosis/resurfacing cycle. (1) Surface TNR epitopes were blocked with Fab fragments directed against TNR (white ellipse), applied together with non-fluorescent anti-mouse nanobodies (dark gray circles). (2) 4 hours later, the newly-emerged TNR epitopes were tagged with new Fab fragments directed against TNR, in the absence of a secondary nanobody. The neurons were then incubated for a further 12 hours, to allow these newly-emerged epitopes to internalize. (3) The newly-emerged TNR epitopes that did not internalize and remained at the surface were blocked with non-fluorescent anti-mouse nanobodies. (4) Immediately afterwards, or 1-3 days later, we revealed the newly-emerged and then internalized TNR epitopes that had again emerged at the surface, by applying STAR635P-conjugated anti-mouse nanobodies (red circles). **e**, The neurons showed very little fluorescence immediately after the second surface blocking step, when imaged in epifluorescence microscopy. The fluorescence intensity increased 1 day later, and even more so 3 days later. This indicates that a sizeable population of TNR molecules has completed a full cycle between the ECM and intracellular compartments during the observation time (n = 3 independent experiments with at least 50 neurons analyzed per experiment, per condition). Statistical significance was evaluated using a Kruskal-Wallis test (H_2_ = 7.2, \**p* = 0.0273, followed by the Dunn multiple comparisons test; \**p* = 0.0199 for the comparison of 0 to 72 h). Scale bar = 10 μm. Data represent the mean ± SEM.

This therefore suggests that TNR molecules endocytose and are then repeatedly recycled. Importantly, this conclusion is independent of any problems with, for example, antibodies leaving their epitopes. Random un-binding of blocking antibodies would allow fluorophore-conjugated TNR antibodies take their place, therefore resulting in immobile, non-dynamic spots on the surface. The un-binding of fluorophore-conjugated TNR antibodies would simply make the respective TNR molecules invisible. Therefore, none of these scenarios would report either endocytosis or recycling of TNR, implying that our interpretation is independent of such problems (which anyway appear to be negligible, Extended Data Fig. 5).

However, a more significant problem is that antibodies may cross-link the TNR molecules, and may thereby change their behavior. To control for this, we repeated our key experiments using monovalent Fab fragments. They could be successfully used for our ‘blocking-labeling’ assay (Extended Data Fig. 6a-d), and showed the recycling of TNR in the same fashion as the antibodies (Extended Data Fig. 6e). The use of Fab fragments also enabled a more elaborate labeling experiment, based on the detection of Fab fragments with unconjugated or fluorophore-conjugated anti-mouse nanobodies (Fig. 3d). This strategy only reveals TNR epitopes that have completed an entire cycle of endocytosis and resurfacing, and again showed ample signals at ∼3 days after the initiation of the experiment (Fig. 3e).

We therefore conclude that a dynamic population of TNR molecules surfaces regularly, preferentially near synapses, and is then endocytosed and recycled over the course of a few days. To verify this conclusion, we blocked dynamin, which is thought to be involved in most endocytosis reactions, and we also perturbed cellular trafficking with monensin or brefeldin. All these treatments perturbed strongly TNR recycling (Extended Data Fig. 7).

To test whether the dynamic pool of TNR is relevant for synaptic transmission, we performed a crude experiment in which these molecules were bound by large aggregates of antibodies ^16^, and we then analyzed synaptic vesicle exo-/endocytosis. This treatment blocked presynaptic activity, while the addition of such aggregates to the other, non-dynamic TNR epitopes, had no effects (Extended Data Fig. 8). We also tested whether our general findings are relevant only for dissociated neuronal cultures. We repeated the ‘blocking-labeling’ assay in cultured hippocampal organotypic slices, where we observed a similar behavior to the dissociated cultures (Extended Data Fig. 9).

Finally, we also sought to extend these observations to other ECM molecules. We used the ‘blocking-labeling’ assay to assess neurocan, chondroitin-sulfate (CS)-bearing proteoglycans (predominantly aggrecan) labeled by *Wisteria floribunda* agglutin (WFA), and hyaluronic acid (HA) ^10^. In a similar fashion to TNR, we observed that the amount of newly-emerged epitopes was far larger than would be predicted from their exceptionally long half-lives ^2,3^, and that the epitope emergence increased after culture activation using bicuculline (Extended Data Fig. 10a-c).

To verify the turnover dynamics of the ECM by a completely different approach, we used a fluorescence recovery after photobleaching (FRAP)-based assay to observe the hyaluronan-binding protein HAPLN1, which is substantially easier to express and monitor than all other molecules tested here (Extended Data Fig. 10d-f, Movie S1). HAPLN1 dynamics were far higher than expected according to its lifetime ^2,3^ and were also significantly faster in synaptic regions, supporting our previous observations for TNR. Moreover, organelle transport of HAPLN1 appeared to take place, as observed in long-term imaging of HALPN1-expressing cultures (Movie S1).

While these last experiments do not constitute a direct proof of endocytosis or recycling for these ECM molecules, they complement the more direct assays used for TNR. Overall, our results demonstrate that the neural ECM is significantly more plastic than previously assumed. In contrast to the belief that the ECM is extremely stable, and that remodeling events are few and far between, we have shown that major constituents of the ECM are in constant transit, and cycle in and out of neurons. This finding is in line with the observation that synapses are constantly changing in the adult brain ^7,8^, even in the absence of discrete plasticity events. As this mechanism may not be limited to TNR, our observations open a new field of investigation that should prove important in understanding not only ECM regulation in the brain, but also brain plasticity and stability in general. Finally, as ECM changes are known to accompany a plethora of brain diseases, these findings should also prove relevant for clinical research in the future.

## Acknowledgements

We thank Dr. Renato Frischknecht from the Leibniz Institute for Neurobiology for providing a mouse deficient in TNR and brevican as well as a control mouse, and Dr. Weilun Sun from DZNE Magdeburg for performing TNR immunostaining and imaging of mouse brain sections. This work was supported by grants to S.O.R. from the German Research Foundation (Deutsche Forschungsgemeinschaft, DFG SFB1190/P09, DFG SFB1286/A03). Also supported by the DFG under Germany’s Excellence Strategy -EXC 2067/1-390729940.

## Author contributions

S.O.R., T.M.D. and A.D. designed the experiments and statistical analysis. T.M.D. performed all fluorescence imaging experiments and the cell-surface biotinylation experiments, with the following exception: the fluorescence imaging of neurocan, hyaluronan and WFA was performed by G.C.P.; P.E.G and H.A.H. assisted with cellular experiments. The cloning of the TNR shRNA AAV was performed by R.K., and the viral infections were performed by S.B. and B.C.. The FRAP experiments were performed by R.K. The electrophysiology experiments were performed by G.B. The data analysis was performed by T.M.D. and S.O.R. The manuscript was written by T.M.D. and S.O.R. and edited by A.D.

## Code and materials availability

all routines are available upon request to T.M.D.

## Methods

### Neuronal cultures

Dissociated primary hippocampal cultures were prepared from newborn rats as previously described ^17,18^. Briefly, hippocampi of newborn Wistar rat pups were dissected in HBSS (140 mM NaCl, 5 mM KCl, 6 mM glucose, 4 mM NaHCO_3_, 0.3 mM Na_2_HPO_4_ and 0.4 mM KH_2_PO_4_) and incubated for one hour in enzyme solution (DMEM containing 0.5 mg/mL cysteine, 100 mM CaCl_2_, 50 mM EDTA and 2.5 U/mL papain, bubbled with carbogen for 10 min). Dissected hippocampi were then incubated for 15 minutes in a deactivating solution (DMEM containing 0.2 mg/mL bovine serum albumin, 0.2 mg/mL trypsin inhibitor and 5% fetal calf serum). The cells were triturated and seeded at a density of ∼ 80,000/cm^3^ on circular glass coverslips (1.8 cm diameter). Before seeding, the coverslips were treated with nitric acid, sterilized and coated overnight with 1 mg/mL poly-L-lysine. The neurons were allowed to adhere to the coverslips for 1-4 hours at 37 °C in plating medium (DMEM containing 3.3 mM glucose, 2 mM glutamine and 10% horse serum), after which they were switched to Neurobasal-A medium (Life Technologies, Carlsbad, CA, USA) containing 2% B27 (Gibco, Thermo Fisher Scientific, USA) supplement, 1% GlutaMax (Gibco, Thermo Fisher Scientific, USA) and 0.2% penicillin/streptomycin mixture (Biozym Scientific, Germany). The cultures were maintained in a cell incubator at 37 °C, and 5% CO_2_ for 14-16 days before use. Percentages represent volume/volume.

For the TNR knockdown experiments, mouse hippocampal cultures were prepared from newborn C57BL/6N mice as previously described (López-Murcia et al., 2019) with minor modifications. For the astrocyte feeder layer, cortices were dissected in HBSS, and enzymatically digested for 15 min at 37°C with 0.05% (w/v) trypsin/EDTA solution (GIBCO). The dissected cortices were then triturated and astrocytes were cultured for 7 days in T-75 flasks in DMEM containing 10% FBS and penicillin/streptomycin (100 U/ml; 100 mg/ml) mixture and 0.1% MITO + serum extender (Biozym Scientific, Germany). After 7 days, the cells were seeded onto 18 mm coverslips at a density of (50,000/cm^3^), and 1 μM FUDR (Sigma) was added to the medium 6 days later. For the neuronal culture, hippocampi were dissected in HBSS, and digested for 1 hour at 37° C in DMEM supplemented with 25 U/ml papain (Worthington Biomedical Corp.), 0.2 mg/ml cysteine (Sigma), 1 mM CaCl_2_, and 0.5 MmM EDTA. Digestion was arrested by a 15 minute incubation in DMEM supplemented with 2.5 mg/ml bovine serum albumin (Sigma), 2.5 mg/ml trypsin inhibitor (Sigma), and 10% (v/v) FBS. Neurons were triturated and seeded onto the astrocyte feeder layers at a density of ∼ 75,000/cm^3^. Prior to the addition of the hippocampal neurons, the culture medium was switched to Neurobasal-A medium (Life Technologies, Carlsbad, CA, USA) containing B27 supplement (Gibco, Thermo Fisher Scientific, USA), 2 mM GlutaMax (Gibco, Thermo Fisher Scientific, USA) and penicillin/streptomycin (100 U/ml; 100 mg/ml) mixture (Biozym Scientific, Germany). At DIV7 the neurons were infected with AAV1/2-GFP-U6-shTNR (shRNA for TNR) or AAV1/2-GFP-U6-shScr (scramble shRNA) (1 μl/ml with a titer of 3.35 × 10^10^). At DIV14, infected neurons were washed extensively in PBS and fixed by immersion in a solution of 4% paraformaldehyde in 0.1 M PBS.

For the FRAP live imaging experiments, rat cortical cultures (from E18-E19 and P0-P3) were prepared as previously described ^19–21^. The neurons were seeded at a density of ∼ 80,000/cm^3^ on circular glass coverslips (1.8 cm diameter), and maintained in Neurobasal medium (Gibco, Thermofisher Scientific, USA), supplemented with B27 (Gibco, Thermofisher Scientific, USA), L-Glutamine (Gibco, Thermofisher Scientific, USA) and penicillin/streptomycin (PAA Laboratories, Pasching, Austria). At DIV7-9, the neurons were infected with the AAV-PSD95-eGFP (1ul per well with a titer of 4.5×10^12^) and AAV-HAPLN1-Scarlet (1μl per well with a titer of 5 × 10^12^). All live imaging experiments were performed on neurons at DIV 21-23 after transferring infected cultures to a Quick Change Chamber 18 mm Low Profile RC-41LP (Warner Instruments), and keeping them in the original culture media at 37°C in the constant presence of humidified carbogen for the duration of the imaging.

Percentages represent volume/volume.

### Preparation and treatment of organotypic hippocampal slice cultures

Organotypic hippocampal slice cultures were prepared as previously described ^22^ with the modifications described in ^23^. In brief, hippocampi of postnatal day 3 (P3) C57BL/6J mice were isolated, and 300-μm thick transverse slices were cut and placed on support membranes (Millicell-CM Inserts, PICMORG50; Millipore). The surface of the slices was covered with culture medium consisting of 50% MEM with Earle’s salts (#M4655; Sigma-Aldrich, Germany), 25 mM HEPES, 6.5 mg/ml glucose, 25% horse serum, 25% Hanks solution buffered with 5 mM Tris and 4 mM NaHCO_3_, pH 7.3. The slices were maintained in a cell incubator at 37 °C and 5% CO_2_ for 14 days before use, and the culture medium was replaced every other day. To block surface epitopes of TNR, the slices were incubated with 50 μg/mL unlabeled Fab fragments directed against TNR (custom-made from the same TNR antibody used in the rest of the work; Synaptic Systems, Göttingen, Germany), applied together with 1 mg/mL unlabeled anti-mouse nanobody, diluted in their own cell media for 2 hours. The slices were subsequently washed in Tyrode’s solution (124 mM NaCl, 30 mM glucose, 25 mM HEPES, 5 mM KCl, 2 mM CaCl_2_, 1 mM MgCl_2_, pH 7.4) and returned to their original conditioned media. For labelling of newly-emerged TNR epitopes, slices were incubated with 2 μg/mL Fab fragments directed against TNR, applied together with 1:500 STAR580-conjugated anti-mouse nanobodies (#N1202-Ab580; NanoTag, Göttingen, Germany). The labelling was performed immediately after blocking or after a further incubation of 12 hours. Throughout the 12 hour incubation, some of the slices were treated with 40 μM bicuculline (#485-49-4; Sigma-Aldrich, Germany) or 0.1% volume/volume DMSO (#67-68-5; Sigma-Aldrich, Germany) as a control.

### Generation of knockdown and expression constructs and production of recombinant adeno-associated particles (AAV)

The shRNA plasmid to knockdown mouse Tenascin-R (GeneID:21960) was cloned by the insertion of the siRNA’s sequence (siRNA ID: SASI_Mm01_00073137, Rosetta Predictions from Sigma Aldrich, Merck) targeting the open reading frame of mouse Tenascin-R into adeno-associated viral (AAV) vector U6 GFP (Cell Biolabs Inc., San Diego, CA 92126, USA), using BamH1 and EcoR1 restriction sites (Okuda et al, 2014). A scrambled siRNA sequence was used as a nontargeting negative control. The sequences for TNR siRNA and control siRNA are 5’-gcgttcactctcctccctg, and 5’-cggctgaaacaagagttgg, respectively. In order to specifically investigate the recovery of hyaluronic acid based matrix around synapses, we designed AAV expression vector carrying mDlg4 (Gene ID: 13385) fused with EGFP with a linker sequence 3xGGGGS in between the ORF to label postsynaptic density, which was custom made as a service by VectorBuilder (Chicago, IL, USA). Furthermore, to label hyaluronic acid-based ECM we designed AAV expression vector carrying link protein hyaluronan and proteoglycan link protein 1 (HAPLN1, Gene ID: 12950) fused with mScarlet subcloned from the plasmid pCytERM_mScarlet_N1 (Addgene plasmid # 85066). Clones were verified by sequencing analysis and used for the production of adeno-associated particles as described previously (Mitlöhner et al, 2020). Briefly, HEK 293T cells were transfected with calcium phosphate method with an equimolar mixture of the expression plasmid, pHelper plasmid, and Rep/Cap plasmid pAAV2/DJ (for the production of the shRNA, pAAV2/1 and pAAV2/2 were combined to have a mixed capsid of AAV1/2). After 48 hours of transfection, cells were lysed using freeze-thaw and treated with benzonase at a final concentration of 50 units/ml for 1 hour at 37°C. The lysate was centrifuged at 8000g at 4°C. The supernatant was collected and filtered with a 0.2-micron filter. Filtered supernatant was passed through pre-equilibrated Hitrap Heparin columns (#17-0406-01; Ge HealthCare Life science), followed by a wash with Wash Buffer 1 (20 mM Tris, 100 mM NaCl, pH 8.0; filtered sterile). Columns were additionally washed with Wash Buffer 2 (20 mM Tris 250 mM NaCl, pH 8.0; filtered sterile). Viral particles were eluted with elution buffer (20 mM Tris 500 mM NaCl, pH 8.0; filtered sterile). Amicon ultra-4 centrifugal filter with a 100,000 molecular weight cutoff were used to exchange the elution buffer with sterile PBS. Finally, viral particles were filtered through 0.22 μm syringe filters (Nalgene®; # Z741696; Sigma-Aldrich, Germany), aliquoted and stored at −80°C until required.

### Immunostaining of TNR KO mouse brain slices

The mice were first deeply anesthetized by ketamine (90 mg/kg of body weight) and xylazine (18 mg/kg of body weight) in a 0.9% NaCl solution and perfused transcardially with 4% paraformaldehyde (PFA) in 0.1 M PBS (PH 7.2) for 10 min. The dissected brains were incubated in 4% PFA containing PBS at 4°C for 2h. Fifty-micrometer thin sagittal sections were cut using a microslicer (HM650V, MICROM). The sections were first washed 3×10min with PBS and then permeabilized with 0.1% Triton X-100 (T9284, Sigma-Aldrich) in PBS for 10 min at room temperature (RT). Next, the sections were incubated for 1 h (at RT with gentle shaking) in a blocking solution containing 10% normal goat serum (16210064, Life Technologies), 0.4% Triton X-100 and 0.1% glycine in PBS. Then the sections were incubated for 24 h (at RT with gentle shaking) with the mixture of *Wisteria floribunda* agglutinin (WFA) (1:500, Vector Laboratories, B-1355) and mouse anti-Tenascin-R (1:200, Synaptic Systems, 400004) for 24h. Then the sections were washed 3x 10 min at RT in PBS and incubated on a shaker for 3 hours at RT with secondary reagents: streptavidin Alexa Fluor® 405 (1:200, Life Technologies (Invitrogen), S32351) and goat anti-mouse Alexa Fluor® 647 (1:200, Life Technologies (Invitrogen), A21236). Afterword, the sections were washed with PBS and then mounted on Superfrost glasses (J1800AMNZ, Thermo Scientific) with Fluoromount medium (F4680, Sigma-Aldrich). Images were acquired with a confocal laser-scanning microscopy (LSM 700, Zeiss) with the same acquisition parameters for the comparison of samples.

### Live-cell immunolabeling and treatments

All treatments and stainings were performed on dissociated hippocampal neurons aged 14 days *in vitro* (DIV14). To block surface epitopes of TNR, neurons were incubated with knock-out-validated antibodies (#217 011; clone 619; Synaptic Systems, Göttingen, Germany) diluted 1:100 in their own cell media, for 2 hours. The neurons were subsequently washed in Tyrode’s solution (124 mM NaCl, 30 mM glucose, 25 mM HEPES, 5 mM KCl, 2 mM CaCl_2_, 1 mM MgCl_2_, pH 7.4) and returned to their original conditioned media. For labeling of newly-emerged TNR epitopes, neurons were incubated with antibodies conjugated to the fluorescent dye Atto647N, Atto550 or STAR580 (custom-made; clone 619; Synaptic Systems, Göttingen, Germany) diluted 1:500 in their own cell media, for 1 hour. The labeling was performed after either 0, 2, 4 or 12 hours post-blocking. After labeling, the neurons were fixed either immediately, or after a further incubation in their original culture media allowing for epitope internalization, as noted in the figure legends. For the experiments with alternative ECM components, surface epitopes were blocked with 1:100 Wisteria floribunda agglutinin WFA (#L8258; Merck, Germany), 1:50 Hyaluronan binding protein HABP (#H0161; Merck, Germany) or 1:100 mouse anti-Neurocan (#N0913; clone 650.24; Merck, Germany) together with 1 mg/mL unlabeled nanobody anti-mouse IgG (custom-made; NanoTag, Göttingen, Germany). Labeling was performed with 1:500 biotinylated WFA or HABP followed by 1:500 streptavidin-Atto647N (#AD 647-61; ATTO-TEC GmbH, Germany), or 1:500 anti-Neurocan and 1:500 nanobody anti-mouse IgG conjugated to STAR635P (#N1202-Ab635P; NanoTag, Göttingen, Germany). Drug applications in these experiments began at the timepoints described below and in the figure legends, and lasted until fixation. To enhance culture activity, neurons were treated with 40 μM bicuculline (#485-49-4; Sigma-Aldrich, Germany) or 0.1% volume/volume DMSO (#67-68-5; Sigma-Aldrich, Germany) as a control, for 12 hours following the blocking step. To reduce culture activity by inhibiting AMPA and NMDA receptors, neurons were treated with 10 μM CNQX (#0190; Tocris Bioscience, Germany) and 50 μM AP5 (#0106; Tocris Bioscience, Germany) for 12 hours following the blocking step. To block the activity of matrix metalloproteinases, neurons were treated with 10 μM GM6001 (#CC1010, Merck, Germany) for 12 hours following the blocking step. To digest glycosaminoglycans, neurons were treated with 0.5 units/mL Chondroitinase ABC from *Proteus vulgaris* (#C3667, Merck, Germany) for 30 minutes following the blocking step. To perturb dynamic-dependent endocytosis, neurons were treated with 30 μM Dyngo^®^ 4a (#ab120689; Abcam, United Kingdom) for 2 hours following the labeling step. To perturb Golgi trafficking, neurons were treated with 5 μg brefeldin (#B7651; Sigma-Aldrich, Germany) or 1 μM monensin (#M5273; Sigma-Aldrich, Germany) for 4 hours, added from the onset of blocking. For the experiments described in Fig. 3d-e and Extended Data Fig. 6, surface TNR epitopes were blocked with 10 μg/mL unlabeled Fab fragments directed against TNR (custom-made from the same TNR antibody used in the rest of the work; Synaptic Systems, Göttingen, Germany), applied together with 1 mg/mL unlabeled anti-mouse nanobody (custom-made; NanoTag, Göttingen, Germany), diluted in their own cell media for 2 hours, and newly-emerged TNR epitopes were labeled with 2 μg/mL Fab fragments directed against TNR, applied together with 1:500 STAR635P or STAR580-conjugated anti-mouse nanobodies (#N1202-Ab635P, #N1202-Ab580; NanoTag, Göttingen, Germany). For the experiment described in Extended Data Fig. 6c-d, newly-emerged epitopes were labeled 12 hours post-blocking. The neurons were mounted for live imaging 4 hours after labeling, and the Fab fragments bound to surface TNR molecules were stripped by incubation with proteinase K (see below). For the experiment described in Extended Data Fig. 6e, newly-emerged TNR epitopes were labeled 4 hours post-blocking. The neurons were mounted for live imaging 1, 2 or 3 days after labeling and the Fab fragments bound to surface TNR molecules were stripped by incubation with proteinase K (see below). For the experiment described in Fig. 3d-e, newly-emerged TNR epitopes were tagged with unlabeled Fab fragments directed against TNR 4 hours later post-blocking. The tagged epitopes that remained exposed on the surface 12 hours later were blocked by an additional incubation with 1 mg/mL unlabeled anti-mouse nanobody for 2 hours. Subsequently, the remaining Fab-tagged epitopes that had internalized were revealed at the surface with 1:500 STAR635P-conjugated anti-mouse nanobodies immediately after the second blocking step, or following an additional incubation of 1-3 days. To visualize the active synaptic vesicle pool, neurons were incubated with 1:500 polyclonal rabbit antibodies directed against the lumenal domain of Syt1 conjugated to the fluorescent dye Oyster488 (#105 103C2; Synaptic Systems, Göttingen, Germany) during the TNR labeling step.

For surface antibody stripping, neurons were incubated with 8 units/ml Proteinase K from *Tritirachium album* (#P2308, Merck, Germany) in Tyrode for 5 min at room temperature. The neurons were then washed and mounted for live imaging (for the experiments described in Fig. 3c and Extended Data Fig. 6c-e), or immediately fixed and post-immunostained (for the experiments described in Fig. 3b and Extended Data Fig. 7a).

Sequestration of new TNR epitopes on the plasma membrane was done as previously described ^16^. To perturb the newly-emerged TNR epitopes, neurons were blocked for 2 hours with unlabeled TNR antibodies (1:100), together with 1 mg/mL unlabeled anti-mouse secondary nanobodies. The neurons were then labeled 12 hours after blocking with TNR antibodies (1:500) together with 1 mg/mL biotinylated secondary nanobodies (custom-made; NanoTag, Göttingen, Germany). This ensures that only the newly-emerged epitopes, which surfaced during the 12 hours after blocking, are tagged by biotin. Alternatively, to perturb specifically the epitopes that do not recycle (“all other epitopes”), the blocking step was performed using unlabeled TNR antibodies (1:100), but without unlabeled anti-mouse secondary nanobodies. We incubated the neurons for 12 hours, to ensure that the recycling epitopes are endocytosed, and we then applied the biotinylated secondary nanobodies. These now cannot detect the endocytosed molecules, and cannot detect the newly-emerged TNR epitopes, since these molecules are not bound by the TNR antibodies. The biotinylated secondary nanobodies only detect the TNR antibodies on the surface molecules that were present at the time of labeling (12 hours previously) and did not endocytose in the meanwhile – meaning the stable, non-recycling pool. Subsequently, neurons were incubated with antibody aggregates diluted 1:20 in Tyrode for 30 min (untreated neurons were incubated in plain Tyrode). The antibody aggregates consisted of goat anti-biotin (#B3640; Merck, Germany) and STAR635P-conjugated donkey anti-goat (#ST635P; Abberior GmbH, Göttingen, Germany). Donkey anti-goat and goat anti-biotin antibodies were diluted 1:10 in PBS, and the mixture was left to rotate overnight at 4°C. Following the addition of antibody aggregates, the neurons were incubated for 5 min with 1:100 polyclonal rabbit antibodies directed against the lumenal domain of Syt1 conjugated to the fluorescent dye Oyster488 (#105 103C2; Synaptic Systems, Göttingen, Germany) in either plain Tyrode (stimulated condition) or Ca^2+^-free Tyrode containing 1mM EGTA and 1 μM TTX (#1069; Tocris Bioscience, Germany) (unstimulated condition). To reveal the releasable vesicles, neurons were stimulated electrically with field pulses with a frequency of 20-Hz and an intensity of 100 mA, for 10 seconds. The stimulations were performed with a 385 Stimulus Isolator and an A310 Accupulser Stimulator (World Precision Instruments, Sarasota, FL, USA), using a custom-made platinum plate field stimulator (8 mm-distanced plates). Following the stimulation, the neurons were incubated a further 5 minutes with the Syt1 antibodies and then briefly washed and fixed. Live-cell incubations were performed at 37°C. Washing steps were performed in pre-warmed Tyrode.

Antibodies were diluted from 1mg/ml stocks, unless specified otherwise.

### Cell-surface biotinylation assay

The assay was adapted from ^13^. Briefly, neurons were incubated with 100 μM Leupeptin (#L2884; Sigma-Aldrich, Germany) for 1 hour at 37°C, to inhibit lysosomal protein degradation. Leupeptin was also present in the cell media throughout the remainder of the experiment. The neurons were incubated with 1.5 mg/mL EZ-Link™ Sulfo-NHS-S-S-Biotin (#21331; Thermo Fisher Scientific, USA) in PBS for 30 minutes at 37°C. The neurons were subsequently washed in PBS containing 10 mM glycine to quench the unreacted biotin. The neurons were either immediately scraped into lysis buffer, to detect the entire surface pool (50 mM Tris-HCl, 150 mM NaCl, 2 mM EDTA, 0.5% IGEPAL, 0.5% sodium deoxycholate, 0125 mM PMSF protease inhibitor, and 1x protease inhibitor cocktail: #87786; Thermo Fisher Scientific, US), or were returned to their original culture media and incubated at 37°C, to allow for endocytosis of cell-surface proteins. After a further 6 hours, the neurons were incubated with glutathione cleavage buffer containing 50 mM glutathione (#G6013; Sigma-Aldrich, Germany), 75 mM NaCl, 10 mM EDTA, 75 mM NaOH and 1% BSA in H_2_O, for 20 minutes at 4°C. Subsequently, the neurons were either returned to their original cell media supplemented with 10 mM glutathione, or quenched in iodoacetamide buffer containing 50 mM iodoacetamide (#A1666; Applichem GmbH, Germany) and 1% BSA in PBS, for 30 minutes at 4°C, and were immediately scraped into lysis buffer, to reveal the endocytosed pool of molecules. The remaining neurons were incubated a further 18 hours at 37°C, to allow for the resurfacing of endocytosed proteins, and were then subjected to a second glutathione cleavage reaction to cleave, the newly-surfacing biotinylated proteins. The neurons were then quenched in iodoacetamide buffer and scraped in lysis buffer, thereby revealing the non-recycling pool.

Biotinylated cell-surface proteins were pulled down with streptavidin-coupled magnetic beads (#11205D; Thermo Fisher Scientific, US). The beads were isolated, were washed, and were then blocked with PBS containing 2.5%, bovine serum albumin (BSA) (A1391-0250; Applichem, Germany) and 0.1 % Tween20 (9005-64-5; Merck, Germany) for 1 hour. They were then immunostained with 1:500 monoclonal mouse anti-TNR, 1:100 monoclonal mouse anti-Syt1 (#105 311; Synaptic Systems, Göttingen, Germany) or 1:100 monoclonal mouse anti-calmodulin (#MA3-917; Thermo Fisher Scientific, US), together with 1:100 STAR635P-conjugated anti-mouse nanobodies, overnight at 4°C in. The beads were subsequently washed and mounted on glass slides in Mowiol for imaging. Confocal imaging was performed on a Leica TCS SP5 microscope (Leica, Wetzlar, Germany) equipped with an HCX Plan Apochromat 63x 1.4 NA oil objective. The 561nm or 633 nm lines of a Helium-Neon laser were utilized for excitation, using acousto-optic tunable filters to select appropriate emission wavelengths. The images were acquired with photomultiplier tubes. For each channel the pinhole was set to 1 Airy unit.

### Electrophysiology

Na^+^, K^+^ currents and mEPSCs were recorded by conventional whole-cell patch-clamping under a physiological temperature of 37°C, with continuous perfusion of ACSF (pre-saturated with 95% O_2_ and 5% CO_2_). ACSF (Artificial cerebrospinal fluid) contains (in mmol/L): 119 NaCl, 2.5 KCl, 2 CaCl_2_, 2 MgSO_4_, 1.25 NaH_2_PO_4_, 26.2 NaHCO_3_, 11 Glucose, with pH ∼7.4 and osmolarity 305-315 mOsm. Patch pipettes (2–4MΩ) were pulled from borosilicate glass (1.6mm outside diameter) and filled with intracellular solution containing (in mmol/L): 130 HMeSO_4_, 5 KCl, 1 EGTA, 10 HEPES, 4 MgCl_2_, 2 ATP-Na_2_, 0.4 GTP-Na, 7 Phosphocreatine-Na_2_, with pH adjusted to 7.2–7.4 by KOH, and osmolarity adjusted to 290–295 mOsm. End concentration (in mmol/L) of K^+^ is ∼135, of Na^+^ is ∼17.4 and of Cl^-^ is 9. Whole-cell configuration was first made under a normal ACSF. Na^+^ and K^+^ currents were elicited by a series of depolarizing pulses between − 60mV and +50mV, in 10mV increments, from a holding potential of − 70mV. mEPSCs were recorded at a holding potential of −70 mV under ACSF, with 1μM TTX and 100μM picrotoxin, after Na^+^ currents have disappeared. An EPC-9 patch-clamp amplifier equipped with Patchmaster software (HEKA Electronics, Germany) was used for data acquisition. No leak currents compensation was used, but fast and slow capacitances ^24,25^, and series resistances were corrected on-line. The data were sampled at 40 kHz and filtered at 10 kHz (four-pole Bessel) and 5.9 kHz (three-pole Bessel). Data were stored and exported to Matlab for further analysis, which was performed by thresholding the curves using an empirically-derived threshold, to detect the mEPPs. Control cultures were exposed to boiled TNR antibodies, while test cultures were incubated with normal TNR antibodies.

### Imaging of fluorescence recovery after photobleaching (FRAP)

FRAP experiments were performed using a Leica SP5 confocal microscope (Leica-Microsystems, Mannheim, Germany; LAS AF software, version 2.0.2) equipped with Argon (458, 476, 488, 496, 514 nm laser lines), Diode Pumped Solid State (DPSS, 561nm) and HeNe (633nm) lasers and acousto-optic tunable filters (AOTF) for selection and intensity adaptation of laser lines. Confocal images with parameters of 512 × 512 pixels display resolution, 8 bit dynamic range, 63× objective, NA 1.40, 3× optical zoom, voxel size approximately 0.16 × 0.16 × 0.3 μm3) were acquired. In all FRAP experiments, one baseline image was collected as prebleach image before 15-16 circular ROIs of 12-15 μm2 size around PSD-95 puncta were bleached by high-intensity 561-nm laser resulting in bleaching of only HAPLN1-scarlet fluorescence. After bleaching, images were acquired at the rate of one image per 10 minutes up to 16-17 hours to monitor the recovery of hyaluronic acid based matrix. Images were acquired with the same settings for all samples before and after bleaching. Data was collected following bleaching at 7 to 16 independent bleaching spots in each experiment. 14 independent experiments (FRAP movies) were performed, using 14 coverslips from 3 independent culture preparations. The images were analyzed by manually selecting circular regions of interest (ROIs) centered on the bleached spots, and monitoring the signal in the ROIs. The signal was corrected for the loss of signal caused by the repeated imaging. This parameter was monitored in ROIs placed in regions from the same regions, which were not bleached initially. The fluorescence signals were normalized to the pre-bleaching intensity, and were then averaged, as shown in the respective figure. The synaptic enrichment values shown in Extended Data Fig. 10f represent the number of synapses identified, normalized to the total PSD95 fluorescence within the respective ROIs.

### Fixation and post-fixation immunostaining of dissociated hippocampal cultures

Neurons were fixed in 4% PFA in PBS (137 mM NaCl, 10 mM Na_2_HPO_4_, 2 mM KH_2_PO_4,_ 2.7 mM KCl, pH 7.4) for 20 min on ice followed by 20 min at room temperature. The fixation reaction was quenched with 100 mM NH_4_Cl in PBS for 30 min. For subsequent immunostainings, neurons were permeabilized and blocked with PBS containing 2.5%, bovine serum albumin (BSA) (A1391-0250; Applichem, Germany) and 0.1 % Tween20 (9005-64-5; Merck, Germany) for 1 hour. To identify excitatory glutamatergic synapses and visualize the synaptic vesicle pool, neurons were immunostained with anti-VGlut1 nanobodies directly conjugated to STAR580 (#N1602; NanoTag, Göttingen, Germany).To identify inhibitory synapses, neurons were incubated with 1:200 rabbit polyclonal anti-VGAT (#131 103; Synaptic Systems, Göttingen, Germany). For the experiment described in Extended Data Fig. 1a, perineuronal nets were revealed by incubating the neurons with 1:500 mouse anti-TNR antibodies (#217 011; clone 619; Synaptic Systems, Göttingen, Germany) or 1:500 biotinylated Wisteria floribunda agglutinin WFA (#L8258; Merck, Germany), diluted in the same blocking solution. The neurons were washed and incubated with 1:200 Cy3-conjugated goat anti-mouse IgG or (#115-165-146; Dianova, Germany) or Cy3-conjugated streptavidin (#016-160-084; Dianova, Germany). Blocking or labeling of surface TNR epitopes in fixed cells was performed identically to these treatments in live cells (see above). To determine the background fluorescence intensity stemming from the unspecific binding of the secondary antibody, the neurons were incubated with 1:500 Atto647N-conjugated goat anti-mouse IgG (#610-156-121S; Rockland, USA) or STAR580-conjugated anti-mouse nanobodies (#N1202-Ab580; NanoTag, Göttingen, Germany). To identify the neuronal axons (for the experiment described in Fig. 3b), neurons were immunostained with 1:100 mouse monoclonal directed against Ankyrin G (#75-146; NeuroMab, USA), followed by incubation with 1:200 Cy3-conjugated goat anti-mouse IgG (#115-035-146; Dianova, Germany). Primary and secondary antibody incubations were performed for 1 hour at room temperature, also in with PBS containing 2.5%, bovine serum albumin (BSA; A1391-0250; Applichem, Germany) and 0.1 % Tween20 (9005-64-5; Merck, Germany). For visualizing neuronal membranes, fixed and immunostained coverslips were incubated with DiO (#D275; Molecular probes, Invitrogen). In brief, DiO crystals were diluted 20 μg/ml in PBS and sonicated for 30 min, and then diluted further to 2 μg/ml. Neurons were incubated with DiO for 20 min at 37°C, washed once and left overnight.

Neurons were subsequently washed twice and embedded in Mowiol (Calbiochem, Billerica, MA, USA).

### Imaging

Two-color STED imaging was performed on an Abberior easy3D STED microscope (Abberior GmbH, Göttingen, Germany) equipped with a UPlanSApo 100×, 1.4 NA oil immersion objective (Olympus Corporation, Shinjuku, Tokyo, Japan) and an EMCCD iXon Ultra camera (Andor, Belfast, Northern Ireland, UK). Pulsed 488 nm, 561 nm and 640 nm lasers were used for excitation, and easy3D module 775 nm laser was used for depletion. For each channel, the pinhole was set to 1 Airy unit. Confocal imaging was performed on a Leica TCS SP5 microscope (Leica, Wetzlar, Germany) equipped with HCX Plan Apochromat 100× and 63x 1.4 NA oil objectives. The 488 nm line of an argon laser, the 561nm and 633 nm lines of a Helium-Neon laser were utilized for excitation, using acousto-optic tunable filters to select appropriate emission wavelengths. The images were acquired with photomultiplier tubes or Hybrid detectors. For each channel the pinhole was set to 1 Airy unit. Epifluorescent imaging was performed on an inverted Nikon Ti microscope (Nikon Corporation, Chiyoda, Tokyo, Japan) equipped with a Plan Apochromat 100×, 1.4 NA oil immersion objective and an IXON X3897 Andor (Belfast, Northern Ireland, UK) camera. For the live imaging experiments described in Fig. 3a and Extended Data Fig. 3, a cage incubator system was fitted (OKOLab, Ottaviano, Italy). The temperature was maintained at 37°C and CO_2_ was maintained at 5%. The neurons were imaged in the own cell media at a rate of 1 frame every 2 hours. For each frame, 5 z-slices were acquired resulting in a stitched mosaic image of 5×5 lateral fields of view. Alternatively, the live imaging experiments described in Fig. 3c and Extended Data Figures 6c-e and 9 were performed on an inverted Olympus microscope (Olympus Corporation, Shinjuku, Tokyo, Japan), equipped with a UPlanFL 20x, 0.5 NA dry objective (Fig. 3c and Extended Data Figures 6e and 9) or a 100× 1.40 NA oil-immersion Plan Apochromat objective (Extended Data Fig. 6c-d) and a charge-coupled device camera (F-View II; Olympus). For the experiments described in Fig. 3c and Extended Data Fig. 6e, five images of different fields of view were acquired before and after treatment with Proteinase K. For the experiment described in Extended Data Fig. 6c-d, an image was acquired in the same field of view before and after treatment with Proteinase K. For the experiment described in Extended Data Fig. 9, images were acquired from three different fields of view for each cultured slice.

### Long-term live imaging

Neurons were plated on 24 well glass-bottomed cell culture plates (#P24-1.5H-N, Cellvis, USA) at a density of ∼50,000 cells per well as described (see main methods). At DIV14, surface TNR molecules were labeled with Atto550-conjugated TNR antibodies, as described above (see <u>Live-cell immunolabeling and treatments section above)</u>. The neurons were transferred to an automated live-cell incubator/imaging system (BioSpa™ 8 Automated Incubator coupled with a Cytation™ 5 Cell Imaging Multi-Mode Reader, BioTek, USA). The plates were incubated in the BioSpa at 37°C and 5% CO_2_ for 4 days. Every 4 hours, the plates were automatically transferred to the Cytation 5, also set to 37°C and 5% CO_2_, and imaged using a 20x Plan Fluorite, 0.45 NA (#1320517, BioTek PN) objective in the RFP imaging channel, in addition to a phase-contrast channel overlay. For each well, 16 fields of view were acquired (a total 6.3 x 4.7 mm imaging area per well).

### Image and data analysis

Analyses were performed in Matlab (MathWorks, Natick, MA, USA) and Python (Python Software Foundation).

#### Analysis of epifluorescent images

Images were first thresholded to remove background signal; regions above an empirically defined threshold were treated to contain real signals for further analysis. For analyzing the epifluorescent images from the experiments described in Figures 2b-c and Extended Data Figures 5a-d, 5g-h, 6a-b, 7b, 8d-e, and 10a-c, regions of interest (ROIs) were manually selected in the images to include neurites, and the mean fluorescence intensity was calculated within the ROIs. For analyzing the epifluorescent images from the experiment described in Extended Data Fig. 2a, ROIs surrounding both neurites and somas were selected. For analyzing the epifluorescent images from the experiments described in Extended Data Figures 1c-d, 5e-f, 7a and 9, the mean fluorescence intensity was calculated for entire images. For the live imaging experiment described in Fig. 3a and Extended Data Fig. 3, the mean intensity was calculated on the maximum intensity projection images acquired at each time point. ROIs were drawn around neuronal somas and the mean intensity for each individual neuron was normalized to the first time point. For the experiments described in Fig. 3c and Extended Data Fig. 6e, ROIs containing neurites were selected on neurites within the images. For each experimental day, a ratio was calculated between the mean intensity before and after incubation with the proteinase. For the experiment described in Extended Data Fig. 6c-d, the images before and after proteinase incubation were aligned, and the mean fluorescence intensity was determined in selected ROIs in corresponding regions of the images.

#### Analysis of confocal images of immunolabeled beads

For analyzing the confocal images of immunolabeled beads (Fig. 1), the images were segmented based on an empirically defined threshold, to detect the individual beads, and the background signal was removed. The total fluorescence intensity was calculated for each bead, and a mean value was calculated per experiment.

#### Analysis of TNR at synapses by STED and confocal microscopy

The synapses in the images shown in Fig. 2d and Extended Data Fig. 4 were identified by manually thresholding the VGlut1 or Syt1 channels., and square regions of interest centered on the individual VGlut1 or Syt1 puncta were excised. For the visualization of synaptic enrichment of TNR, as depicted in Fig. 2d, the excised image segments were automatically rotated to maximally overlap in the DiO channel, and averaged to one single image illustrating TNR localization at an average synapse (following procedures previously described ^26,27^). For the enrichment analysis at dendrites and axons, the mean TNR fluorescence intensity was calculated for all image segments in concentric circles of increasing radii. To quantify the amount of newly-emerged TNR epitopes after 12 hours specifically at synapses, as shown in Fig. 2e, the mean fluorescence intensity of TNR was estimated in VGlut1-positive pixels. For the correlation of TNR fluorescence to the presynaptic recycling vesicle pool size, as depicted in Extended Data Fig. 4, the image segments surrounding individual presynapses (determined by the Syt1 channel) were sorted by the mean Syt1 fluorescence intensity, and then binned in five ordinal groups, to include a similar number of synapses. For each experiment, the mean fluorescence was calculated in each bin, and was normalized to the median intensity of the respective experiment. A similar procedure was used to analyze the TNR fluorescence in relation to the postsynaptic spines: image segments were excised for synaptic regions based on the VGlut1 channel, and mushroom-shaped spines were identified manually in these segments through the DiO channel. The spine size (area) was determined by thresholding the DiO signal of each individual image segment and counting the pixels above the threshold. The image segments were sorted by the spine size, and then binned in five ordinal groups, to include a similar number of synapses. For each experiment, the mean fluorescence and size were calculated in each bin, and normalized to the median of the respective experiment.

#### Analysis of TNR at neurites and synapses by confocal microscopy

For the comparison of TNR in axons and dendrites as depicted in Fig. 3b, ROIs were selected on the axons and dendrites separately, based on the DiO channel and the AnkyrinG staining, and the mean TNR fluorescence was calculated within these ROIs. For colocalizing TNR with the excitatory and inhibitory synapse markers VGlut1 and VGAT as depicted in Extended Data Fig. 1b, an image-wide pixel-by-pixel correlation was first calculated between the images. The pixels indicating a high correlation between each of the channels were selected, and the TNR amount corresponding to these pixels was calculated as % of the total TNR staining in the images (corrected for background).

### Statistical analysis

Statistical significance was calculated with two-tailed t-tests, Mann-Whitney U-tests, ANOVA, Kruskal-Wallis or Friedman tests, and post-hoc analyses were calculated with Bonferroni, Holm Sidak, Fisher’s LSD or Dunn tests, as noted in the figure legends. For repeated measures ANOVA, the Greenhouse-Geisser adjustment was applied to account for any departures from sphericity.Correlations were calculated with Pearson’s R or Spearman’s ρ. A p-value of < 0.05 was considered statistically significant.

## Extended Data Figures

**Extended Data Fig. 1.**
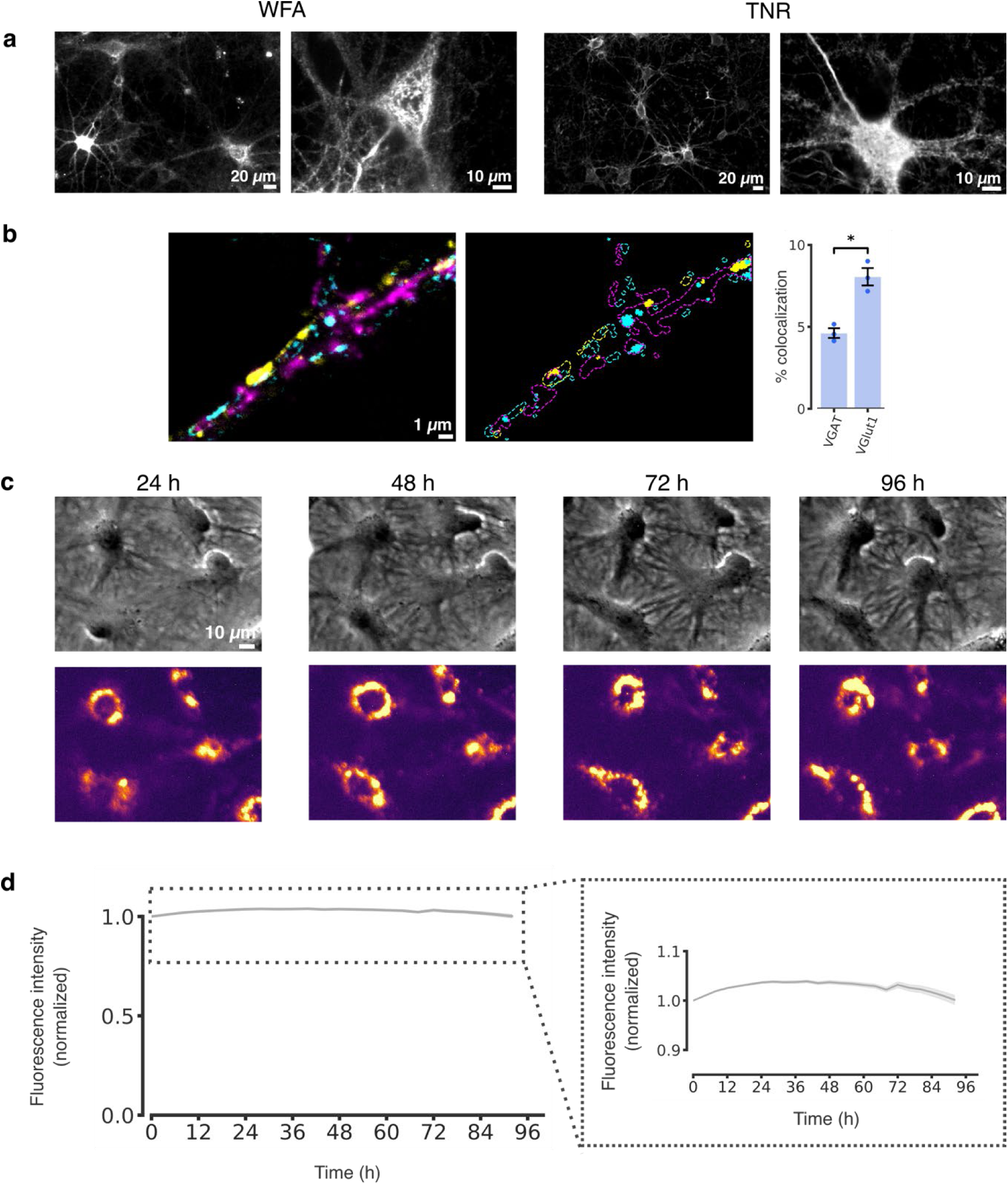
Antibody-labeled TNR molecules are present in the perineuronal net and at synapses, and are extremely stable. **a**, We fixed neuronal cultures at 14 days *in vitro* (DIV14), and we then labeled them with *Wisteria floribunda* agglutin (WFA), which binds chondroitin-sulfate (CS)-bearing proteoglycans ^28^ (top panels), or with TNR antibodies (bottom panels), and imaged them with epifluorescence microscopy. Both labels exhibited lattice-like structures that surrounded the somas and proximal dendrites of a subset of neurons (perineuronal nets; PNNs), which also suggests that the ECM has reached a sufficient level of maturity in these cultures. Scale bar = 20 μm (left panels), 10 μm (right panels). **b**, To assess the amount of TNR at excitatory and inhibitory presynaptic boutons, we labeled all TNR epitopes by incubating live neurons with fluorophore-conjugated TNR antibodies (magenta). Glutamate-releasing (excitatory) and GABA-releasing (inhibitory) boutons were determined by immunostaining the synaptic vesicle markers VGlut1 (yellow) and VGAT (cyan), respectively. The right panel shows the boundaries of the boutons and of TNR domains. The regions where high intensities of TNR and synaptic signals overlap are indicated by full shading in yellow or cyan. TNR, VGlut1 and VGAT are imaged in confocal microscopy. Scale bar: 1 μm. The colocalization of TNR with VGlut1- and VGAT-positive presynapses was determined by measuring the amount of TNR in fully colocalizing pixels, as a percentage of the total TNR staining in the images. A higher proportion of all TNR epitopes can be found at excitatory synapses (n = 3 experiments, at least 10 neurons imaged per experiment; paired *t*-test: t = 4.684, \**p* = 0.043). Data represent the mean ± SEM, with dots indicating individual experiments. **c**, Surface TNR molecules in live neurons were labeled with fluorophore-conjugated TNR antibodies, as in panel a. We then imaged the neurons for up to 96 hours using an automated cell incubator/microscope setup (BioSpa Live Cell Analysis System, BioTek, USA). Top panels: phase-contrast images of a single field of view after 1, 2, 3 and 4 days. Bottom panels: the corresponding images in the fluorescent channel. Scale bar = 10 μm. **d**, The graph shows the mean fluorescence intensity, analyzed over the entire images, normalized to the t_0_ timepoint. A very small overall loss of signal is seen, confirming the slow degradation and turnover of TNR, as known from the literature (n = 3 experiments, >100 neurons per experiment). Data represent the mean (line) ± SEM (shaded region).

**Extended Data Fig. 2.**
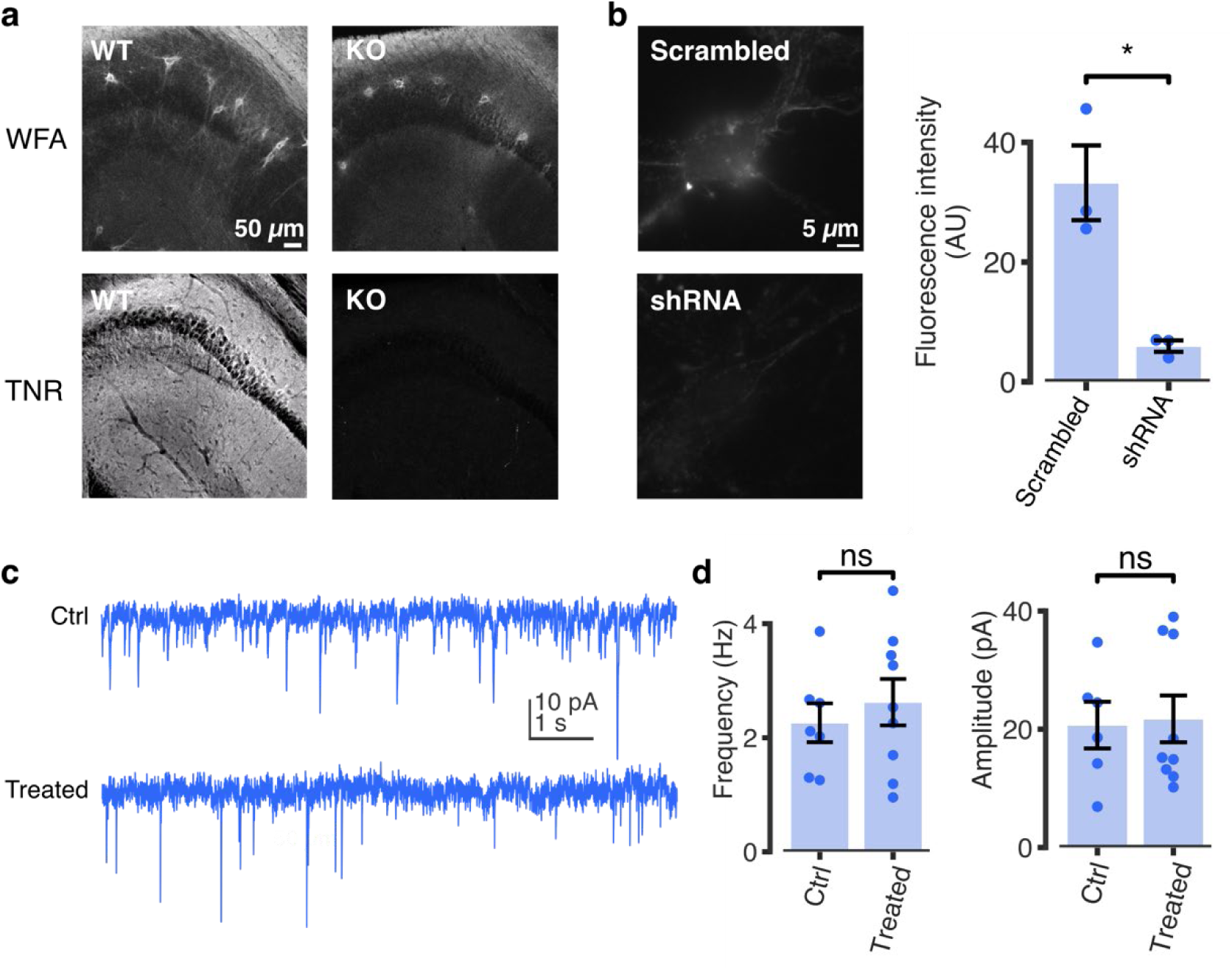
Validation of TNR antibodies. **a-b**, The TNR antibodies used throughout this study were validated in both knockout and knockdown neurons. **a**, Hippocampal slices of Brevican/TNR KO hippocampal slices, imaged in epifluorescence microscopy. Top panels: *Wisteria floribunda* agglutin (WFA) staining. Bottom panels: TNR staining. The TNR signal is entirely lost in the KO. Hippocampal CA2 regions are shown. Scale bar = 50 μm. (C) **b**, Neuronal cultures were infected at DIV7 with AAV vectors co-expressing eGFP together with shRNA against TNR, or a scrambled control. Sample images of neurons fixed at DIV21 from scrambled-(top panel) or shRNA-treated cultures (bottom panel). Scale bar = 5 μm. An analysis of the mean fluorescence intensity (from epifluorescence microscopy experiments) indicates that TNR is significantly reduced in KD cultures, to ∼ 20% of the amount in untreated cultures. Statistical significance was evaluated using a Student’s *t*-test (n = 3 individual experiments, t = 4.562, \**p* = 0.045). The slow turnover of TNR, with a half-life of ∼7 days in rat neuronal cultures (Dörrbaum et al., 2018), suggests that ∼25% of the TNR should still be present after a knockdown treatment of two weeks. At the same time, this experiment was performed in mouse cultures, for which the AAV vectors were optimized, while other measurements from this work, and from the literature, refer to rat cultures. This implies that turnover values may be somewhat different in this experiment, which was performed solely to verify the antibodies used here. **c-d**, Treatment with TNR antibodies does not affect synaptic transmission. **c**, We measured miniature EPSCs in control rat hippocampal cultures (top) and in cultures treated with TNR antibodies (bottom). Scale bar = 10 pA (vertical axis) and 1 s (horizontal axis). **d**, We analyzed the mEPSC frequency and amplitude in the cultures, and found no significant differences (7 independent experiments for control, 9 for cultures treated with TNR antibodies). Statistical significance was evaluated using Mann-Whitney U tests (U = 28, *p* = 0.758; U = 27, *p* > 0.999 for the comparison of frequency and amplitude, respectively). Data represent the mean ± SEM, with dots indicating individual experiments.

**Extended Data Fig. 3.**
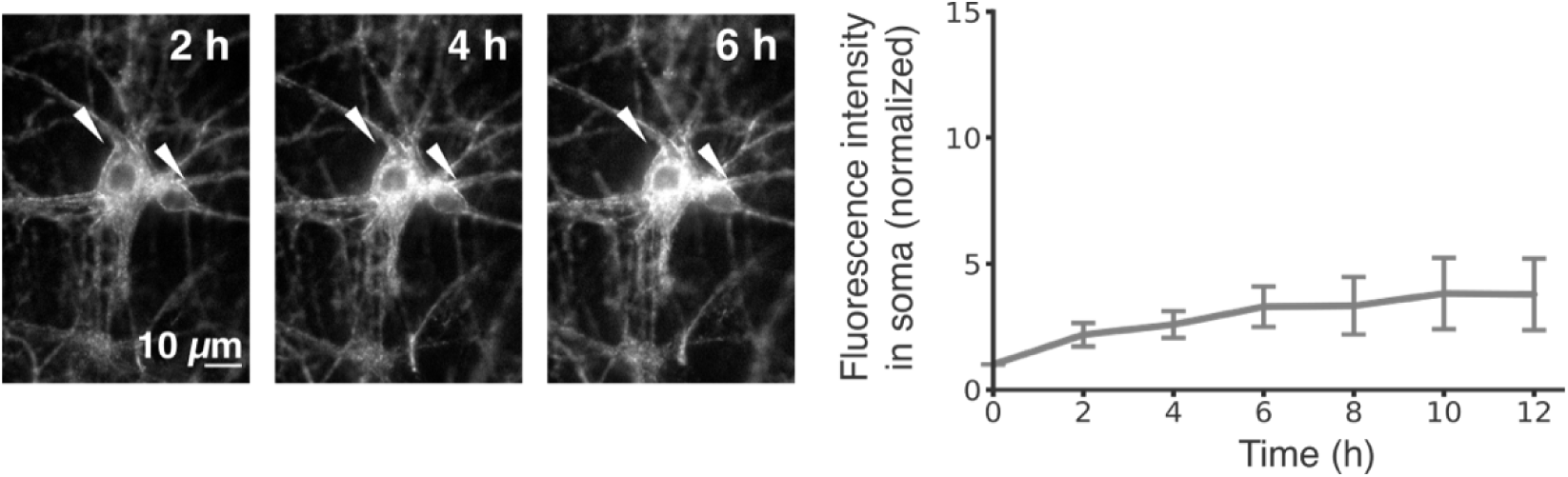
TNR molecules endocytose over several hours. Surface TNR molecules were labeled with fluorophore-conjugated TNR antibodies, and were monitored by live epifluorescence imaging over 12 hours. TNR molecules accumulate in the soma over time, but many do not appear to change their location, and neurites remain fully visible over time. Arrowheads indicate cell somas. Scale bar = 10 μm. The plot shows the mean fluorescence intensity in the cell somas, normalized to the t_0_ timepoint. A significant increase in the signal over time confirms the suggestion that some proportion of the TNR molecules are endocytosed (n = 5 independent experiments, Mann-Whitney U test between the t0 timepoint and the all sequential timepoints: U = 25, \**p* = 0.016). Data represent the mean ± SEM.

**Extended Data Fig. 4.**
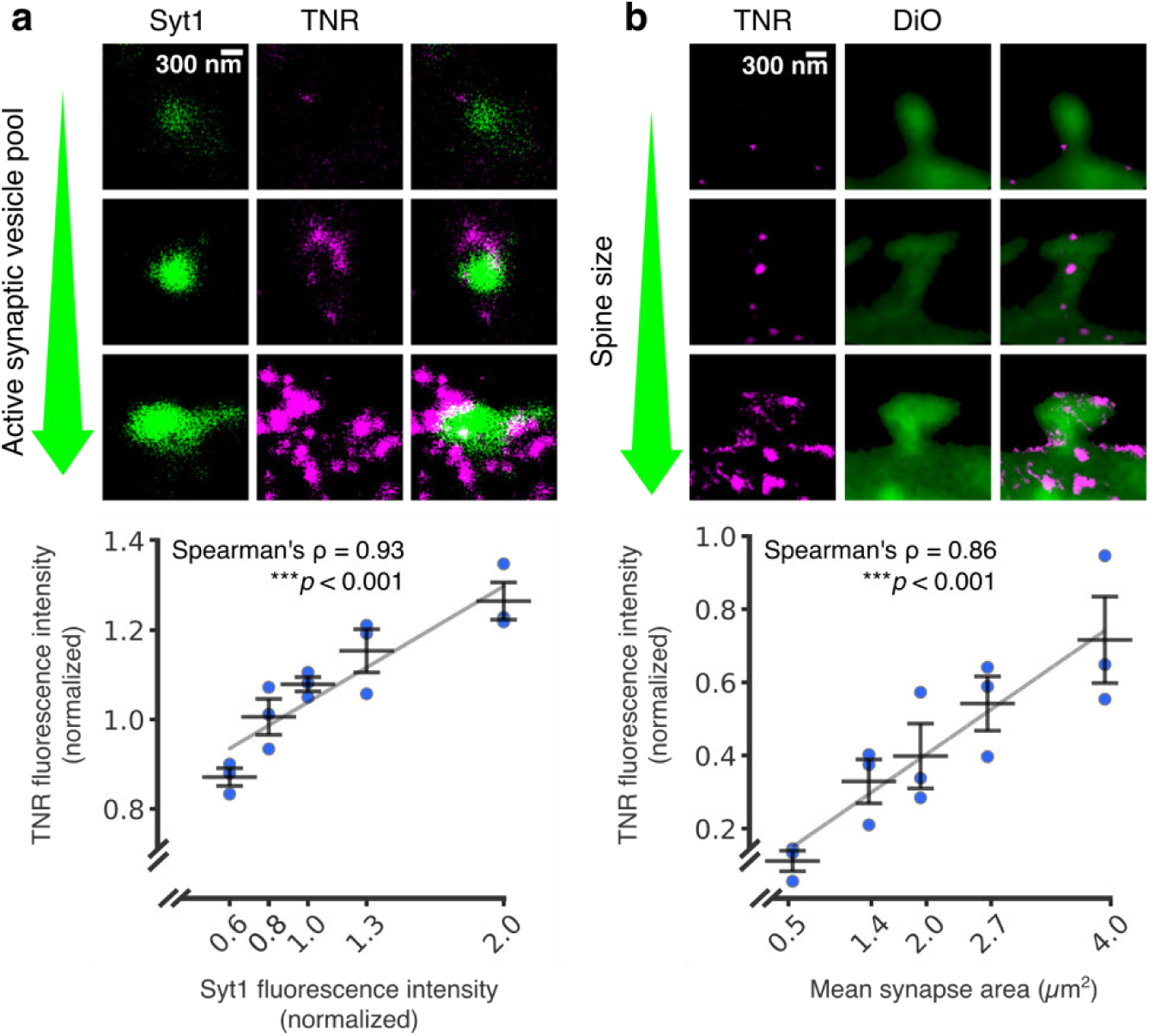
The emergence of TNR epitopes is dependent on synaptic weight. **a**, The TNR epitopes in the ECM were blocked as in the previous experiments, and 12 hours later the cultures were incubated with fluorophore-conjugated TNR antibodies (magenta) and with fluorophore-conjugated antibodies for the intra-vesicular domain of Syt1 (green), which reveal the synaptic vesicle pool that undergoes exo- and endocytosis (the active pool). The size of this pool is a measure of the activity of the respective boutons. The panels show example synapses with different active vesicle pools, imaged in STED (TNR) and confocal (Syt1). Scale bar = 300 nm. The graph shows the mean fluorescence intensities normalized to the median intensity of the respective experiment. The Syt1 intensities are binned to include an equal number of synapse images. An analysis of the correlation of the TNR signal at Syt1-labeled synapses indicates that the TNR signals correlate strongly with the size of the active vesicle pool in the respective boutons (n = 3 independent experiments, > 1100 synapses per data point, Spearman’s ρ = 0.927, \*\*\**p* < 0.001). **b**, Newly-emerged TNR epitopes (magenta) were labeled after 12 hours as in panel a, and the neuronal plasma membrane was visualized with DiO (green). The panels show example spines with different sizes, imaged in STED (TNR) and confocal (DiO). Scale bar = 300 nm. The graph shows the mean fluorescence intensity of TNR and the mean synapse area, normalized to the median values in the respective experiment. The synapse area values are binned to include an equal number of synapse images. An analysis of the correlation of the TNR signal at DiO-labeled spines indicates that the TNR signals correlate strongly with the size of the dendritic spine for newly-emerged TNR epitopes (n = 3 independent experiments, > 280 synapses per data point, Spearman’s Rho = 0.862, \*\*\**p* < 0.001). All data represent the mean ± SEM, with dots indicating individual experiments.

**Extended Data Fig. 5.**
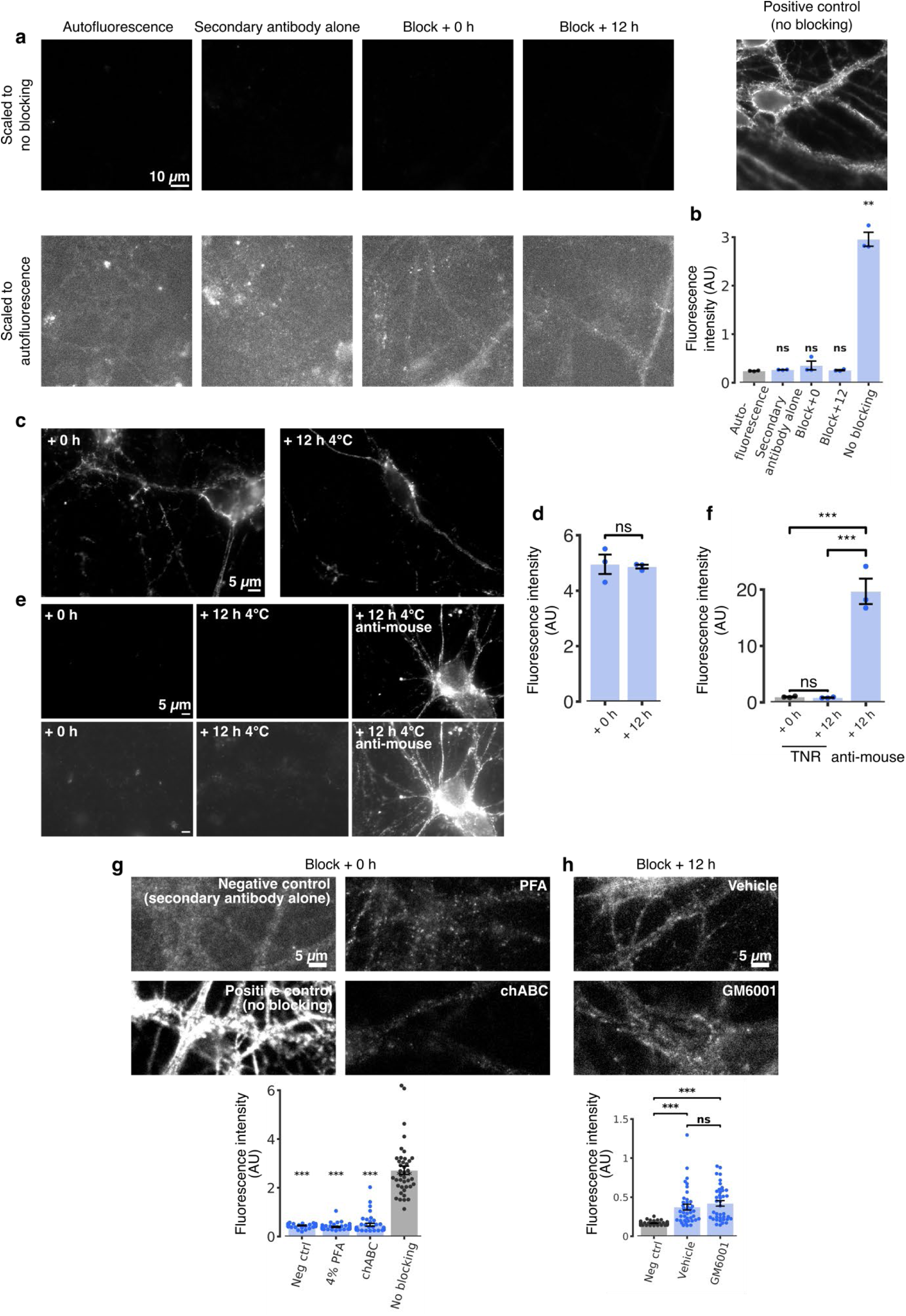
Validation of the antibody assay for revealing newly-emerged TNR epitopes. **a-b**, The TNR antibodies do not separate from their epitopes in fixed cells at 37 °C. **a**, We fixed neuronal cultures, and we then blocked their surface TNR epitopes with non-fluorescent antibodies. Immediately afterwards, or after 12 hours, we incubated the neurons with Atto647N-conjugated TNR antibodies. As a control, all TNR epitopes were labeled, by omitting the blocking step. The blocked cultures showed virtually no detectable fluorescence when imaged in epifluorescence microscopy (top panels). In the bottom panels, we enhanced the image contrast, to reveal the outlines of the cells. These are as bright in these images as in negative controls exposed only to Atto647N-conjugated secondary anti-mouse antibodies (leftmost panel). Scale bar = 10 μm. **b**, The analysis of the mean fluorescence intensity confirms that no new TNR epitopes emerge after 12 hours of incubation in fixed neurons, indicating that the blocking antibodies persist on their epitopes, and do not allow the Atto647N-conjugated TNR antibodies to bind (n = 3 independent experiments, at least 10 neurons imaged per data point). Statistical significance was evaluated using ANOVA (F_4, 10_ = 120.3, \*\*\**p* < 0.001), followed by Holm-Sidak multiple comparisons test (\*\*\**p* < 0.001 for the comparison of each mean to the ‘no blocking’ condition). None of the other conditions were significantly different from the autofluorescence negative control. All data represent the mean ± SEM, with dots indicating individual experiments. **c-f**, The TNR antibodies do not separate from their epitopes in live cells at 4 °C. **c**, We labeled live neuronal cultures with Atto647N-conjugated TNR antibodies, and imaged them either immediately or following a 12 hour-long incubation at 4°C, in epifluorescence microscopy. Scale bar = 5 μm. **d**, An analysis of the mean fluorescence intensity shows that no significant change in staining is apparent after 12 hours at 4°C, indicating that the antibodies persist on their epitopes (n = 3 independent experiments, at least 10 neurons imaged per data point). Statistical significance was evaluated using a paired *t*-test (t = 0.286, *p* = 0.802). **e**, We blocked surface epitopes with unlabeled TNR antibodies, and fixed them either immediately or following a 12 hour-long incubation at 4°C. We then labeled the neurons with Atto647N-conjugated TNR antibodies, to reveal the available TNR epitopes, made available by the putative un-binding of blocking TNR antibodies, or with Atto647N-conjugated mouse secondary antibodies, to reveal the unlabeled blocking TNR antibodies. We then imaged the neurons in epifluorescence microscopy. Scale bar = 5 μm. **f**, An analysis of the fluorescence intensity shows that the staining with TNR antibodies is similar, and extremely low, both immediately and 12 hours after the blocking step (n = 3 independent experiments, at least 10 neurons imaged per data point). Statistical significance was evaluated using one-way ANOVA (F_2. 6_ = 69.32, *p*\*\*\* < 0.001) followed by Holm-Sidak multiple comparisons test (*p* = 0.945 for the comparison between ‘TNR 0 h’ and ‘TNR 12 h’, and *p*\*\*\* < 0.001 for both comparisons to the ‘anti-mouse’ condition). All data represent the mean ± SEM, with dots indicating individual experiments. **g-h**, The labeled TNRs are not pre-existing extracellular epitopes or the result of cleavage of existing ECM structures. **g**, To test whether the newly-emerged epitopes observed after TNR blocking could represent pre-existing epitopes that simply become available to antibody binding, we blocked surface epitopes with unlabeled antibodies, and then subjected them to several treatments that severely modify the cell surface. The treatments were fixation with 4% PFA or digestion of the surface glycosaminoglycans with chondroitinase ABC. We then added fluorophore-conjugated TNR antibodies to assess the number of epitopes that become available through these procedures. As positive and negative controls we used neurons incubated with TNR antibodies where we omitted the blocking step, or neurons incubated with anti-mouse secondary antibodies, respectively. We then imaged the neurons in epifluorescence microscopy. Scale bar = 10 μm. The graph (bottom panel) indicates that all conditions show substantially less fluorescence than the positive control, and are not distinguishable from the negative control (n = 40 neurons for each condition, from 2 individual experiments for ‘chABC’ and ‘No blocking’ conditions, and from 3 individual experiments for the ‘neg ctrl’ and ‘PFA-fixed’ conditions). Statistical significance was evaluated using a Kruskal-Wallis test (H_3_ = 89.06, \*\*\**p* < 0.001), followed by the Dunn multiple comparisons test for comparing each mean to the ‘no blocking’ condition (\*\*\**p* < 0.001). **g**, To test whether the newly-emerged TNRs could represent new epitopes that are exposed through the cleavage of existing ECM structures by secreted proteases, we blocked surface epitopes with unlabeled antibodies, and then treated the cultures with GM6001 to block the activity of matrix metalloproteinases (or with DMSO, as a control). We then added fluorophore-conjugated TNR antibodies to assess the amount of epitopes that become available, and imaged the neurons with epifluorescence microscopy. Scale bar = 10 μm. The graph (bottom panel) indicates that drug-treated cultures do not differ significantly from a negative control where neurons were incubated with anti-mouse secondary antibodies (n = 40 neurons for each condition). Statistical significance was evaluated using a Kruskal-Wallis test (H_2_ = 53.34, \*\*\**p* < 0.001), followed by the Dunn multiple comparisons test for comparing the ‘neg ctrl’ condition to ‘vehicle’ and ‘GM6001’(\*\*\**p* < 0.001), and ‘vehicle’ to ‘GM6001’ *(p =* 0.95). All data represent the mean ± SEM.

**Extended Data Fig. 6.**
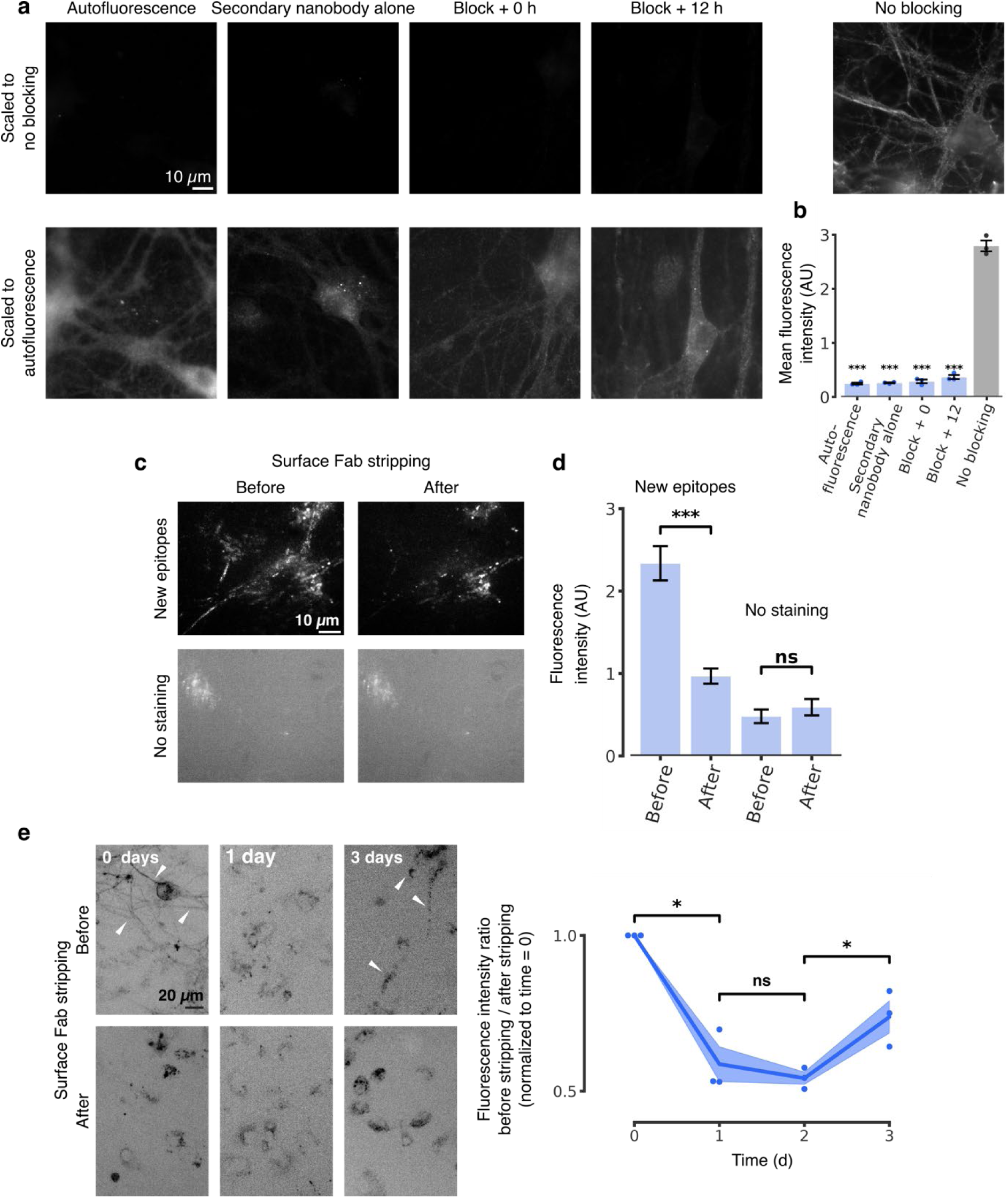
The experiments with antibodies are validated with Fab fragments directed against TNR. **a**, We fixed neuronal cultures, and we then blocked their surface TNR epitopes with Fab fragments directed against TNR, applied together with non-fluorescent anti-mouse nanobodies. Immediately afterwards, or after 12 hours, we incubated the neurons with Fab fragments directed against TNR, applied together with STAR580-conjugated anti-mouse nanobodies. As a control, all TNR epitopes were labeled, by omitting the blocking step. The blocked neurons showed virtually no detectable fluorescence when imaged in epifluorescence microscopy (top panels). In the bottom panels, we enhanced the image contrast, to reveal the outlines of the cells. These are as bright in these images as in negative controls exposed only to STAR580-conjugated secondary anti-mouse nanobodies (leftmost panel). Scale bar = 10 μm. **b**, The application of Fab fragments and STAR580-conjugated anti-mouse nanobodies after 12 hours of incubation in fixed neurons results in no discernible fluorescence signal, indicating that the blocking Fab fragments (coupled to non-fluorescent anti-mouse nanobodies) persist on their epitopes (n = 3 independent experiments, at least 10 neurons imaged per data point). Statistical significance was evaluated using one-way ANOVA (F_4,10_ = 1.191, \*\*\**p* < 0.001), followed by Holm-Sidak multiple comparisons test (\*\*\**p* < 0.001 for the comparison of each mean to the ‘no blocking’ condition). None of the other conditions was significantly different from the autofluorescence negative control. All data represent the mean ± SEM, with dots indicating individual experiments. **c**, We blocked surface TNR epitopes with Fab fragments directed against TNR, applied together with non-fluorescent anti-mouse nanobodies. Newly-emerged TNR epitopes were labeled 12 hours later with new Fab fragments directed against TNR, applied together with STAR635P-conjugated anti-mouse nanobodies. The Fab fragments bound to surface TNR molecules were then stripped by incubation with proteinase K after a further incubation period of 4 hours. The samples were imaged in epifluorescence microscopy before and after the antibody stripping, and were compared to unstained neurons. Scale bar = 10 μm. **d**, A quantification of the mean fluorescence intensity indicates that a significant amount of surface TNR molecules were stripped at 4 hours after staining, whereas no such reduction was observed for unstained neurons. The amount of intracellular molecules that persisted after stripping was higher than background levels of fluorescence, indicating that a portion of the TNR molecules has endocytosed (n = 28 and 42 regions of interest analyzed from 2 sets of images for the ‘new epitopes’ and ‘no staining’ conditions respectively). Statistical significance was evaluated using two-way mixed ANOVA (F_1, 136_ = 36.58, \*\*\**p* < 0.001 for the interaction staining x time) followed by Sidak multiple comparisons test (\*\*\**p* < 0.001 and *p =* 0.727 for ‘before’ vs. ‘after’, for stained and unstained neurons, respectively). All data represent the mean ± SEM. **e**, TNR epitopes labeled with Fab fragments recycled back to the plasma membrane. This experiment reproduced the assay presented in Fig. 3c, but using Fab fragments instead of antibodies. We blocked surface TNR epitopes with Fab fragments directed against TNR, applied together with non-fluorescent anti-mouse nanobodies, and labeled the newly-emerged TNR epitopes 4 hours later with new Fab fragments directed against TNR, applied together with fluorophore-conjugated anti-mouse nanobodies. We then measured the fraction present on the surface after different time intervals. To determine this, we imaged the neurons (in epifluorescence) before and after stripping the surface molecules using proteinase K. At day 0 (immediately after labeling), the stripping procedure strongly reduced the staining. The effect was less visible at 1 day after staining, but became again evident at 3 days after staining, indicating that a high proportion of TNR molecules returned to the surface at 3 days after labeling. Scale bar = 20 μm. We quantified this by reporting the fluorescence ratio between the images taken before and after stripping (normalized to the day 0 time point). The amount stripped at day 3 is significantly higher than at days 1 and 2 (n = 3 individual experiments). Scale bar = 20 μm. Statistical significance was evaluated using repeated measures one-way ANOVA (F_1.519, 3.037_ = 40.91, \*\**p* = 0.007), followed by Fisher’s LSD test (‘day 0’ vs. ‘day 1’: \**p* = 0.018; ‘day 1’ vs. ‘day 2’: *p* = 0.522; ‘day 2’ vs. ‘day 3’: \**p* = 0.044). Data show mean (lines) ± SEM (shaded regions), with dots indicating individual experiments. All measurements in this figure refer to neurites, to avoid the bias caused by the higher autofluorescence of the cell bodies, which is evident in such epifluorescence images.

**Extended Data Fig. 7.**
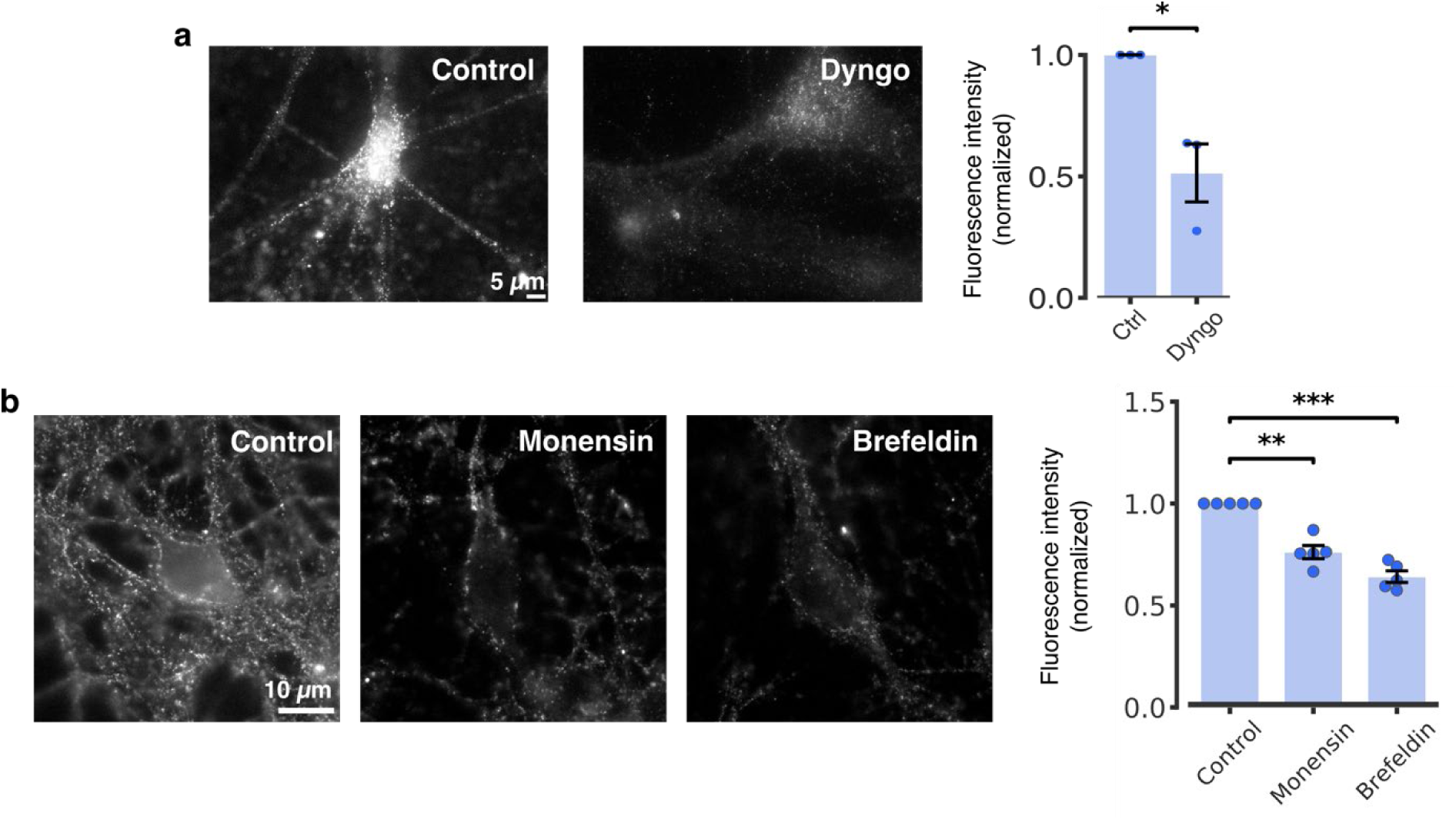
Perturbing endocytosis and cellular trafficking reduces TNR recycling. **a**, Newly-emerged TNR epitopes were labeled 12 hours post-blocking, by the application of fluorophore-conjugated antibodies for 1 hour. Immediately after labeling, dynamin-mediated endocytosis was blocked by incubating the neurons with 30 μM Dyngo^®^ 4a for 2 hours, after which the neurons were stripped by incubation with proteinase K, to reveal only the intracellular epitopes, and imaged with epifluorescence microscopy. The images were compared to control neurons treated with DMSO. Scale bar = 5 μm. The graph shows the mean fluorescence intensity normalized to the control condition in the respective experiment. The drug treatment significantly reduced the amount the internalized TNR epitopes. This treatment cannot be expected to inhibit endocytosis completely in this assay, as the antibodies need to be applied for one hour before the drug, which enables a significant level of endocytosis before the drug can take effect, and because Dyngo at this concentration is not expected to completely remove dynamin function. Statistical significance was evaluated using a paired *t*-test (n = 3 independent experiments, at least 10 neurons imaged per datapoint, t = 4.076, \**p*= 0.015). **b**, The emergence of new TNR molecules was inhibited by drugs perturbing ER/Golgi traffic. We imaged newly-emerged TNR epitopes (in epifluorescence microscopy) after treatments of 7 hours with DMSO (as a control), with the Golgi-stressing ionophore monensin ^29,30^, or with the COPI-disturbing inhibitor brefeldin ^31^. Scale bar = 10 μm. The graph shows the mean fluorescence intensity normalized to the control condition in the respective experiment. Both drugs reduced substantially the amount of newly-emerged TNR epitopes. Statistical significance was evaluated using paired *t*-tests with Bonferroni correction for multiple comparisons (n = 4 independent experiments for each condition, at least 10 neurons imaged per datapoint, t = 7.359, \*\**p* = 0.004 and t = 12.61,\*\*\**p* < 0.001 for the ‘monensin’ and ‘brefeldin’ conditions respectively). Data represent the mean ± SEM, with dots indicating individual experiments.

**Extended Data Fig. 8.**
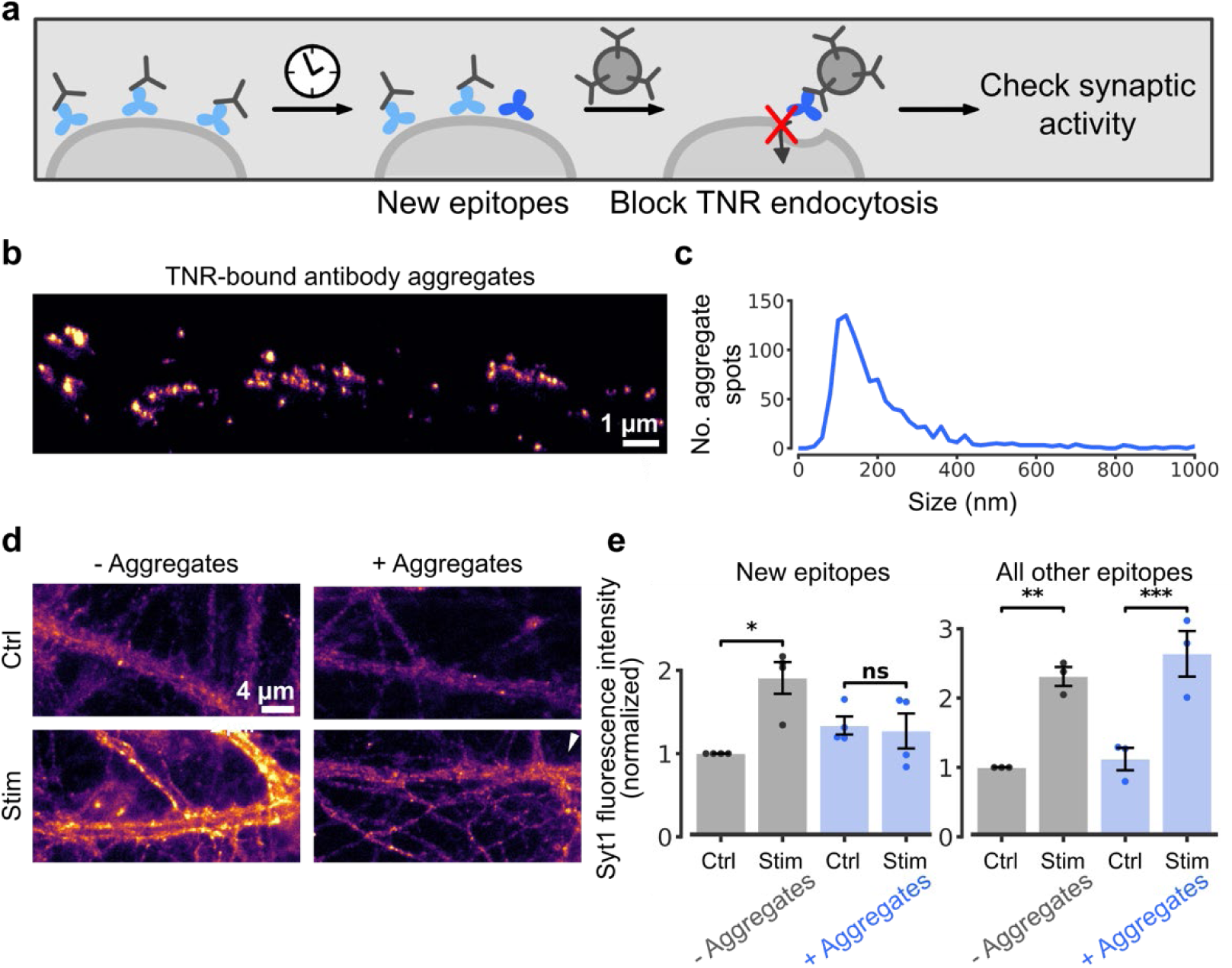
Perturbing the recycling pool of TNR molecules modulates synaptic function. **a**, To test the effects of TNR perturbation on synaptic activity, we labeled the newly-emerged TNR epitopes 12 hours post-blocking with biotinylated antibodies, which were then bound to large antibody aggregates consisting of goat anti-biotin and donkey anti-goat antibodies. As a control, we labeled only those epitopes that had persisted at the cell surface for 12 hours (all other epitopes, meaning non-recycling epitopes). **b**, The aggregates were imaged by STED microscopy. Scale bar = 1 μm. **c**, The size of the antibody aggregates identified in the images was determined by their full width at half maximum (FWHM). The histogram shows the number of aggregate spots at increasing sizes (995 spots were analyzed in total). The highest frequency of spots has a size of ∼120 nm. **d**, We then tested synaptic activity, relying on Syt1 antibodies, as used in Extended Data Fig. 4. Axonal branches (imaged in epifluorescence) are depicted either before stimulation (control, top panels), or after a 10-second, 20-Hz stimulus, which causes the exo-and endocytosis of 50% of the active vesicle pool (the other ∼50% are already present on the surface membrane, waiting for endocytosis ^15,32^). In the absence of stimulation, the Syt1 antibodies detect the surface vesicle population (40-50% of actively-recycling vesicles ^15^). After stimulation the signal increases through the exo- and endocytosis of new vesicles. This is evident for the controls, but not for the aggregate-treated cultures. Scale bar = 4 μm. **e**, The graphs show the mean Syt1 fluorescence intensity, normalized to the untreated, non-stimulated control of the corresponding experiment. An analysis of the intensity confirms that stimulation enhances exo-/endocytosis in control samples, but not in aggregate-treated samples. In contrast, when only the surface-resident, non-recycling TNR epitopes (“all other epitopes”) are bound to antibody aggregates, no difference is observed in the Syt1 signal intensity between control and aggregate-treated cultures. Statistical significance was evaluated using two-way repeated-measures ANOVA on rank-transformed data (F_1, 6_ = 12.54, \**p* = 0.012 (‘new epitopes’); F_1, 4_ = 1.5, *p* = 0.288 (‘all other epitopes’) for the interaction Stim/ctrl x +/-Aggregates) followed by Sidak multiple comparison test (\**p* = 0.02 and *p* = 0.419 (‘new epitopes’); \*\**p* = 0.002 and\*\*\**p* < 0.001 (‘all other epitopes’) for ‘stim’ vs. ‘ctrl’ for untreated and treated neurons, respectively. N = 4 (‘new epitopes’) and 3 (‘all other’) experiments per condition, at least 15 neurons imaged per data point. Data represent the mean ± SEM, with dots indicating individual experiments.

**Extended Data Fig. 9.**
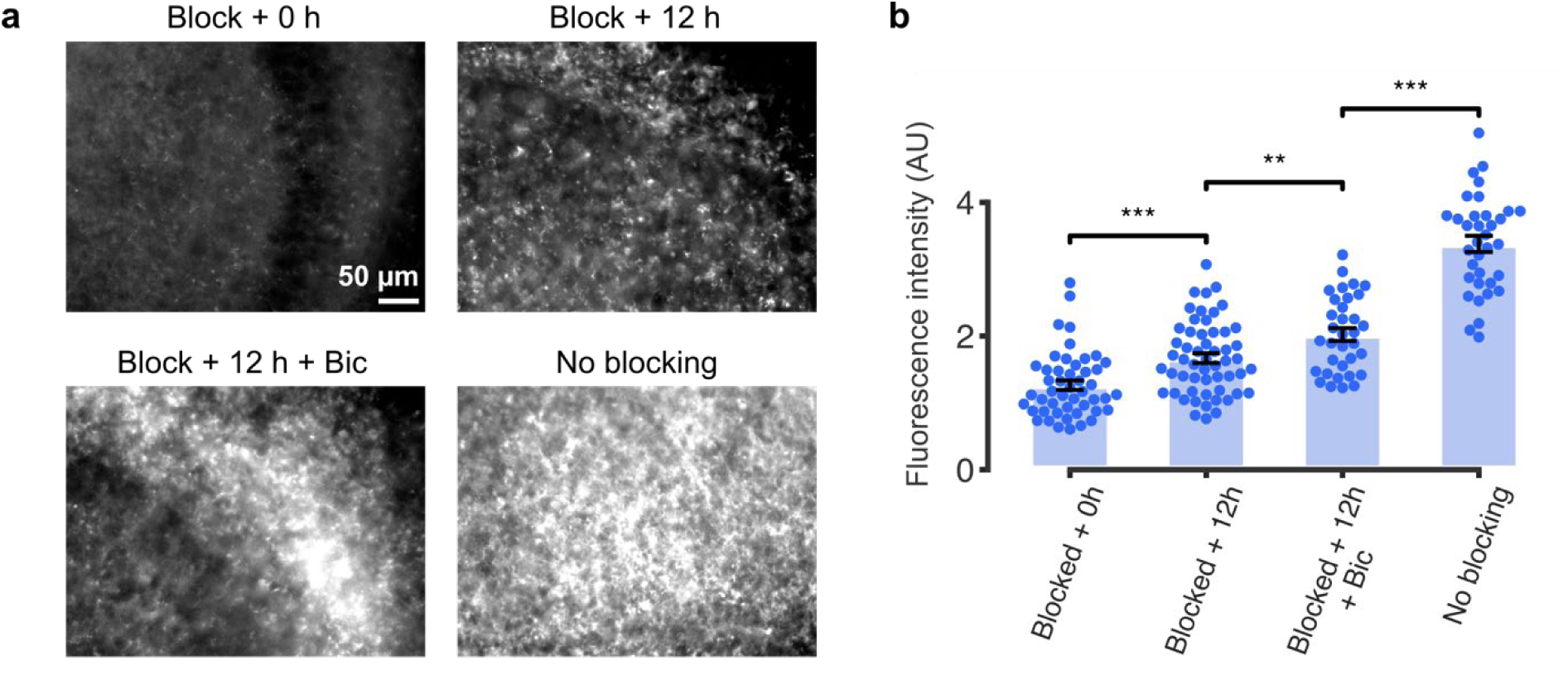
TNR dynamics can also be observed in cultured slices. **a**, To test whether Fab fragments can be used to block TNR surface epitopes in a model that is closer to the *in vivo* morphology of neuronal tissue, we applied our ‘blocking-labeling’ assay to hippocampal organotypic slice cultures. We first blocked TNR surface epitopes by incubating live slices with Fab fragments directed against TNR, applied together with non-fluorescent anti-mouse nanobodies, for 2 hours. Newly-emerged TNR epitopes were labeled with new Fab fragments directed against TNR, applied together with STAR580-conjugated anti-mouse nanobodies, 12 hours after blocking. We compared control cultures with cultures treated with bicuculline (40 μM). To test the efficacy of the blocking procedure, we also labeled the newly-emerged TNR epitopes immediately after the blocking step. The blocked slices showed little fluorescence when imaged in epifluorescence microscopy, when compared with a full surface labeling, in the absence of blocking. Scale bar = 50 μm. **b**, An analysis of the intensity confirms that little TNR staining is visible immediately after blocking. Newly-emerged epitopes can be detected 12 hours post-blocking, and their amounts are higher in the presence of bicuculline (n = 15, 10, 9 and 9 images from 4, 5, 3 and 3 experiments, for ‘blocked + 0 h’, ‘blocked + 12 h’, ‘blocked + 12 h + bic’ and ‘no blocking’ conditions, respectively). Statistical significance was evaluated using one-way ANOVA (F_3, 168_ = 100, \*\*\**p* < 0.001), followed by Holm-Sidak multiple comparisons test (\*\**p* < 0.001 for the comparison of ‘blocked + 0h’ to ‘blocked + 12h’, and ‘blocked + 12 + bic’ to ‘no blocking’; \*\**p* = 0.005 for the comparison of ‘blocked + 12h’ and ‘blocked + 12h + bic’). Data represent the mean ± SEM.

**Extended Data Fig. 10.**
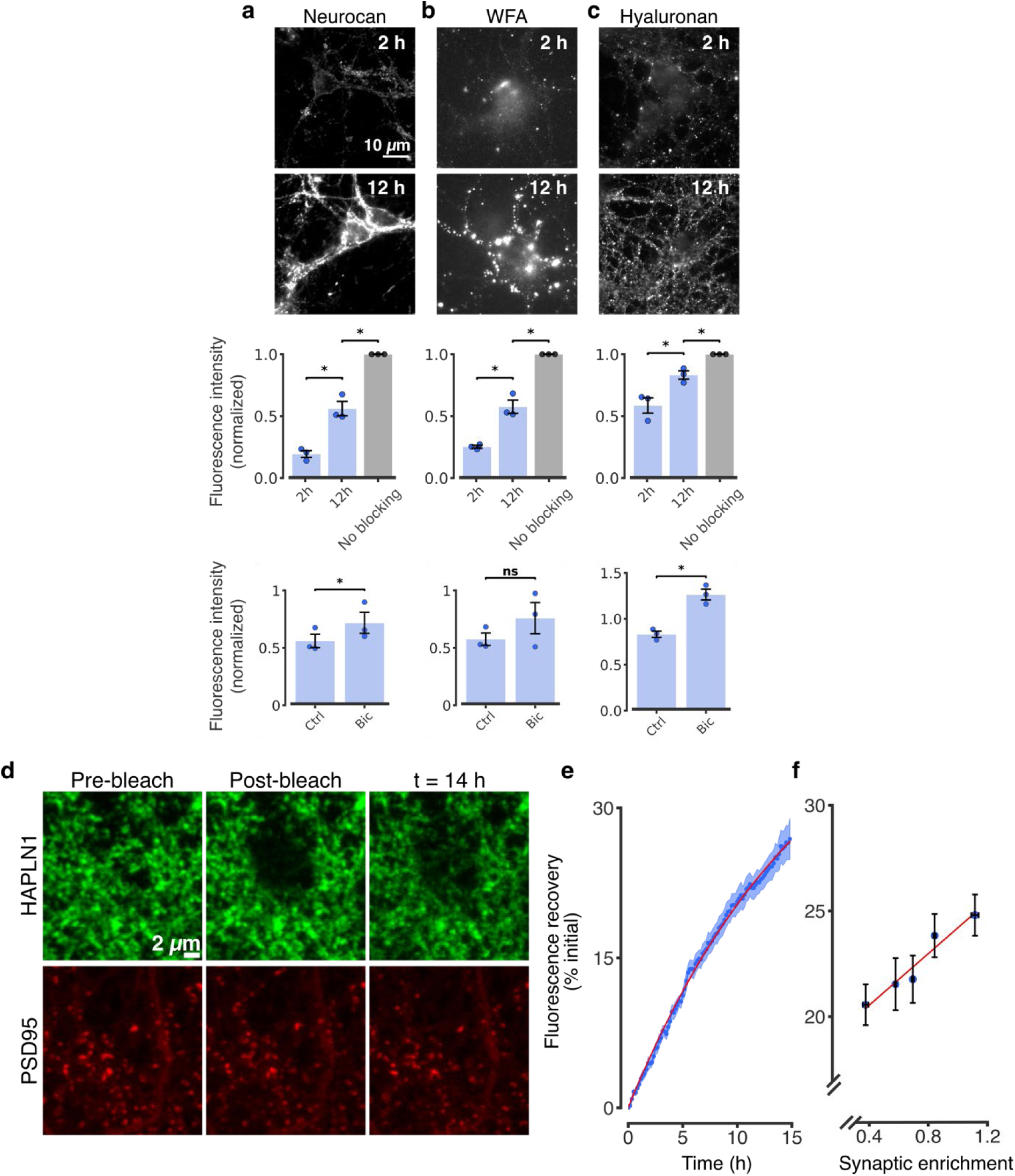
Additional ECM molecules are more dynamic than previously presumed, and surface in neurons in an activity-dependent fashion. **a-c**, Top row: experiments were performed as in Fig. 2b using antibodies for neurocan (panel a) and hyaluronan (panel c), or with *Wisteria floribunda* agglutin (WFA), which labels chondroitin-sulfate-bearing proteoglycans (panel b). The cultures were then imaged in epifluorescence microscopy. Scale bar = 10 μm. Middle row: The graphs show the mean fluorescence intensity normalized to a control condition in which the blocking step was omitted. A gradual increase in newly-emerged epitopes was measured, similarly to the dynamics observed for TNR (n = 3 independent experiments for each condition, at least 10 neurons imaged per experiment; repeated measures one-way ANOVA: F_1.005, 2.011_ = 85.74,\**p* = 0.011; F_1.013, 2.026_ = 119.8, \*\**p =* 0.008; F_1.095, 2.190_ = 42.81, \**p* = 0.018, followed by Fisher’s LSD test: \**p =* 0.050 and \**p* = 0.017; \**p =* 0.036 and \**p* = 0.016; \**p =* 0.034 and \**p* = 0.038 for the comparisons between ‘2 h’, and ‘12 h’ and ‘12 h and ‘no blocking’ conditions for neurocan, WFA and hyaluronan respectively). Bottom row: we compared control cultures with cultures in which network activity was enhanced by inhibiting GABA_A_ receptors using bicuculline (40 μM). The graphs show the mean fluorescence intensity normalized to the mean of the bicuculline-treated condition. The bicuculline treatment had a significant effect for 3 molecules (n = 3 individual experiments, at least 10 neurons imaged per data point; neurocan: t = 4.57, \**p* = 0.045; WFA: t = 1.925, *p* = 0.194; hyaluronan: t = 4.661, \**p* = 0.043). Data represent the mean ± SEM, with dots indicating individual experiments. **d-f**, A FRAP-based assay also demonstrates fast ECM dynamics at synapses. **d**, We used FRAP to observe the local turnover dynamics of the hyaluronan-binding protein HAPLN1. The postsynaptic marker PSD95 (red) and HAPLN1 (green) were expressed in neurons, as fluorescent protein chimeras (see Methods). At time 0, regions of ∼5 μm radius were bleached in the HAPLN1 channel, using a strong laser pulse. The cells were then imaged once every 10 minutes, for 14 hours, in confocal microscopy. Scale bar = 2 μm. **e**, The fluorescence recovery was analyzed, and fitted well to a single exponential process (red curve), which provided a half-life of approximately 12 hours for the recovery process. The curve shows mean (lines) ± SEM (shaded regions) from 14 independent experiments. **f**, To determine whether turnover dynamics differ in synaptic regions, the density of synapses was calculated for the analyzed areas. A strong correlation was found between the percent of fluorescence recovery and synaptic enrichment, indicating that turnover is significantly higher in synaptic regions (R^2^ = 0.941, \*\**p* = 0.006).

**Supplementary Movie 1 FRAP-based assay to observe the local turnover dynamics of the hyaluronan-binding protein HAPLN1**. Time-lapse video of HAPLN1 (green) and PSD95 (magenta), imaged once every 10 minutes, for 14 hours. Organelle-like movement is evident in both channels, indicating ongoing trafficking of proteins. Scale bar = 3 μm.

